# TGFβ signaling systems are prone to inhibition and ligand competition by coreceptor

**DOI:** 10.64898/2026.03.01.708909

**Authors:** Wisam A. Fares, Kevin A. Janes

**Affiliations:** Department of Biomedical Engineering, University of Virginia, Charlottesville, VA 22908, U.S.A; Department of Biochemistry & Molecular Genetics, University of Virginia, Charlottesville, VA 22908, U.S.A

**Keywords:** TGFBR3, betaglycan, ENG, endoglin, GDF11, growth differentiation factor 11

## Abstract

Transforming growth factor-β (TGFβ) ligands and receptors interact with overlapping selectivity to form different signaling complexes. This many-to-many wiring provides versatility in how ligands are perceived and cell types become activated, but the systems-level impact of coreceptors, which bind TGFβ ligands yet do not signal, remains unresolved. We examined the role of canonical surface-bound coreceptors by numerically simulating TGFβ ligand-receptor systems of increasing complexity and pairing results with single-cell measurements in acutely stimulated cells. Using sampled combinations of biologically plausible rate parameters and initial conditions, we find that coreceptors inhibit downstream signaling six times more often than they promote it, and the models reconcile this inhibitory bias with the prevailing view that coreceptors either promote or inhibit signaling. In multi-ligand systems, coreceptor inhibition causes stimuli to be perceived differently by altering competition, and this threshold is confirmed in cells engineered to induce a single coreceptor heterogeneously. Coreceptors are among the most variably expressed components of TGFβ signaling, suggesting that cells exploit coreceptors as a way to switch active ligand-receptor landscapes with a single gene.

## INTRODUCTION

Ligands and receptors of the transforming growth factor β superfamily (collectively, TGFβ) are important signaling molecules in development, homeostasis, and disease.^1^ Canonical TGFβ signaling initiates with ligand binding to a type II receptor (R_II_), followed by heterotrimerization with a type I receptor (R_I_), which becomes activated and then phosphorylates R-SMAD transcription factors that alter gene expression (Figure S1A).^2^ There are seven R_I_s, five R_II_s, and over 30 TGFβ ligands that combine with overlapping selectivity to yield various heterotrimer combinations.^1-3^ Mathematical modeling suggests this competitive binding architecture confers sophisticated modes of ligand perception, cellular addressing, and context-dependent signal transduction.^4-10^

TGFβ signaling complexity is further increased by coreceptors, which bind ligands with high affinity on the cell surface but do not directly signal.^11^ Instead, coreceptors modulate surface access of ligands to R_I_ or R_II_, and the structurally related coreceptors TGFBR3 and ENG do so through bivalent interactions with the TGFβ ligand dimer.^12^ TGFβ coreceptors defy straightforward assignments of function—they may promote or inhibit depending on the definition of signaling (R_I_ or R_II_ complex formation, SMAD phosphorylation, gene expression, growth inhibition) and the cellular context. Quantitative rules of thumb are lacking but important because of the different settings in which TGFβ coreceptors are dynamically regulated,^13-16^ as we confirm here with public datasets.^17-19^

To a minimal model of TGFβ signaling,^4^ we add a second path from ligand to R_II_ through coreceptor and find that prior analytical approaches to signaling output are inaccurate or impossible. We substitute numerical solutions of steady state heterotrimer as the signaling readout then deeply sample plausible ranges of all interaction affinities and protein abundances including coreceptor (∼10^7^ models in total). When these initial conditions are traversed as a function of coreceptor abundance, the models predict that coreceptor is predominantly inhibitory for signaling. Rarer and more-complex dose-response relationships with coreceptor are enhanced in models that encode multiple ligands or receptors yielding heterotrimers that signal with different potency. In engineered cells expressing a single inducible coreceptor, we find support for inhibitory and promote-then-inhibit trends with coreceptor when measuring proportional R-SMAD phosphorylation by multicolor flow cytometry. Models of competitive multi-ligand, multi-receptor systems further predict that coreceptor accentuates asymmetric heterotrimer signaling in ways that trigger altered ligand perception when coreceptor is moderately elevated, as confirmed by experiment. Our combined results suggest that entire TGFβ signaling landscapes are reconfigurable by the abundance of a single coreceptor, which has implications for the apparent pleiotropy of many TGFβ ligands.^1^

## RESULTS

### *TGFBR3* and *ENG* are among the most variably abundant TGFβ (co)receptor genes

To gauge the biological significance of abundance changes in R_I_, R_II_, and coreceptor, we extracted measures of transcript variation in normal and diseased tissues from public datasets. TGFβ ligands are important developmental morphogens that elicit dose-dependent phenotypes depending on cell type.^20^ We assessed bulk variation in TGFβ (co)receptors by quantifying their spread among adult tissues profiled by the Genotype-Tissue Expression Consortium.^17^ Germline effects on overall expression were accounted for by measuring the interquartile range (IQR) from 10 or more tissues per individual and then summarizing IQR values for each gene across hundreds of donors (Figures 1A and S1B), By this measure, most TGFβ receptors were relatively stable, with IQR values for log_2_ transcripts per million (TPM)+1 not discernibly greater than two (i.e., fourfold interquartile variation). *TGFBR3* and *ENG* both exceeded this threshold and were matched by only one R_II_, *TGFBR2*, which is susceptible to extensive epigenetic regulation.^21^ Although TGFBR3 is historically ascribed to endocrine or reproductive tissues and ENG to the endothelium, they are detected in other cell types at different setpoints that may be functionally important.^22,23^

**Figure 1.**
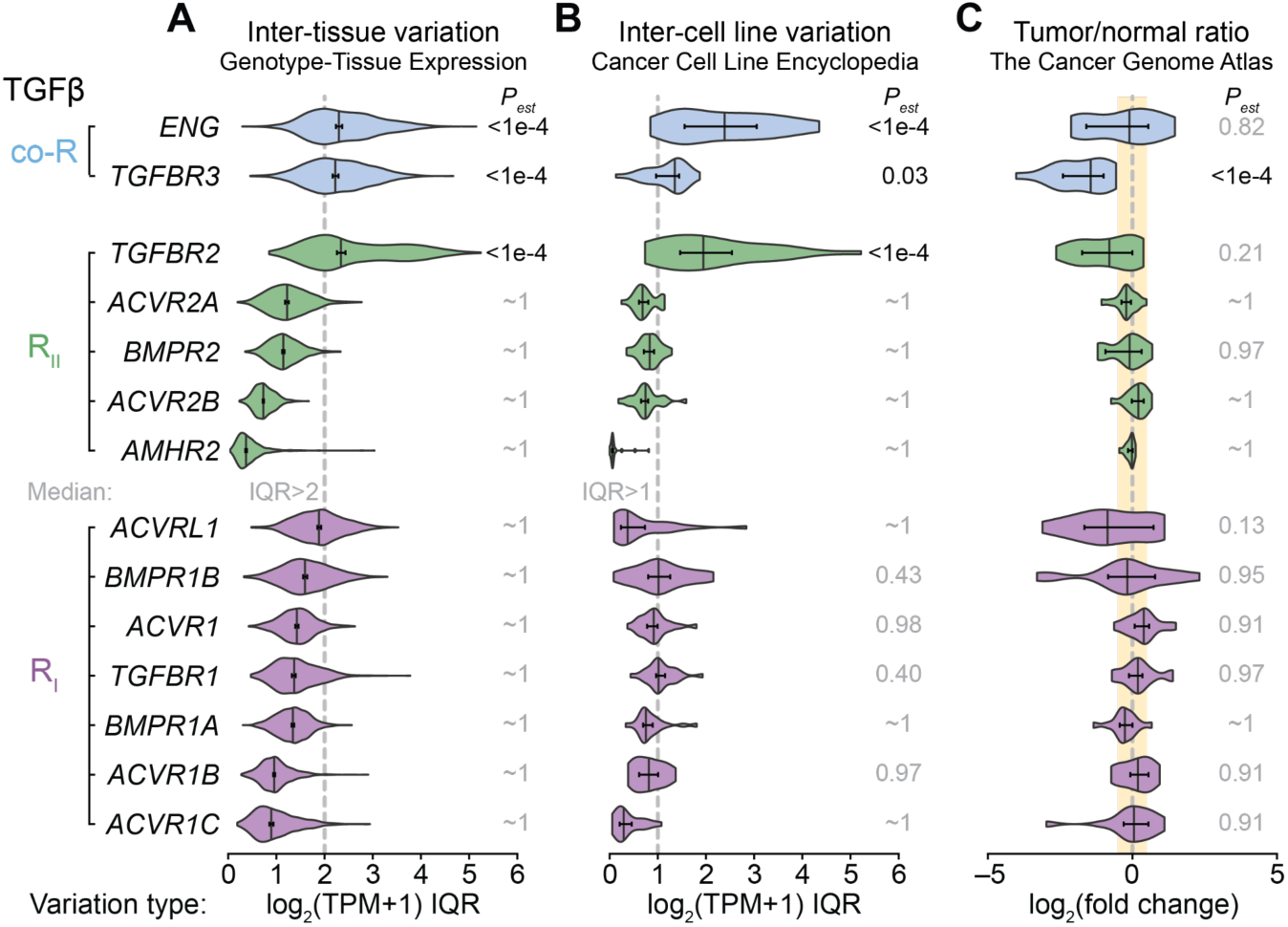
TGFβ coreceptor abundance varies considerably among human tissues, cell lines, and disease states. (A and B) Interquartile range (IQR) of transcript abundance for the indicated TGFβ coreceptor (co-R), type II receptor (R_II_), and type I receptor (R_I_) measured as log_2_ (transcripts per million [TPM] + 1) across (A) 11–39 tissues available from *N* = 830 human donors in the Genotype-Tissue Expression portal^17^ and (B) 8–188 cell lines derived from *N* = 24 cancer types in the Cancer Cell Line Encyclopedia.^18^ (C) Median log_2_ fold-change estimates of tumor-vs.-normal transcript abundance based on TPM from *N* = 15 cancer types each with 13–109 paired tumor-normal samples in The Cancer Genome Atlas.^19^ Violin plots show the median IQR or median log_2_ fold change (vertical line), 95% bootstrapped confidence interval of the median, and estimated *P* value (*P_est_*) for genes with median log_2_ IQR greater than 2 (A) or 1 (B) or absolute log_2_ fold change greater than 0.5 (C) (dashed band). See also Figure S1.

Separately, we asked whether (co)receptors were variably altered in cancer, a disease in which TGFβ misregulation is known to occur.^1^ Using the Cancer Cell Line Encyclopedia,^18^ expression IQR was calculated among cell lines within a given cancer type and then aggregated across two dozen cancer types for each gene (Figures 1B and S1C). We found that cell-line variation within cancer types was rather limited: *TGFBR2*, *ENG*, and *TGFBR3* were the only (co)receptors with an interquartile variation greater than twofold. To address whether the observed deviations related to cell transformation, we compared tumor–normal expression ratios for 15 cancer types (comprising nearly 700 paired samples) in The Cancer Genome Atlas (Figures 1C and S1D).^19^ Only TGFBR3 was appreciably outside of an absolute log_2_ fold change (|log_2_FC|) of 0.5, consistent with its frequent classification as a tumor suppressor.^22^ We conclude that transcriptional regulation of TGFβ coreceptor abundance is as divergent as the most variably expressed TGFβ receptor (*TGFBR2*), meriting an analysis of how coreceptors impact TGFβ signaling.

### Extending a minimal model of TGFβ signaling with coreceptor

Membrane-proximal signaling from TGFβ ligands is canonically modeled as a series of bimolecular binding events that culminate in an active heterotrimer.^4,24-27^ We partially adopted the naming convention of Antebi et al.^4^ and began with one TGFβ ligand (“L”), one R_II_ (“A”), and one R_I_ (“B”) (Figure 2A). The ligand binds R_II_ with a forward rate that is assumed constant for all extracellular:membrane binding events (*k_fe_*). The resulting AL heterodimer (“D” in Antebi et al.^4^) dissociates with rate *k_rD_* or binds to R_I_ with a forward rate parameter that is assumed constant for all membrane:membrane binding events (*k_fm_*). We retained the convention that R_I_ cannot bind ligand without R_II_, which is an accurate approximation for TGFβ and related ligands.^2^ The surface enhancement of 2D binding interactions is captured by setting *k_fm_* >> *k_fe_* (STAR Methods). The resulting ALB heterotrimer (“T” in Antebi et al.^4^) likewise dissociates with rate proportional to *k_rT_*, and the steady state concentration of ALB is taken as the signaling output for a given configuration of rate parameters and initial conditions.

**Figure 2.**
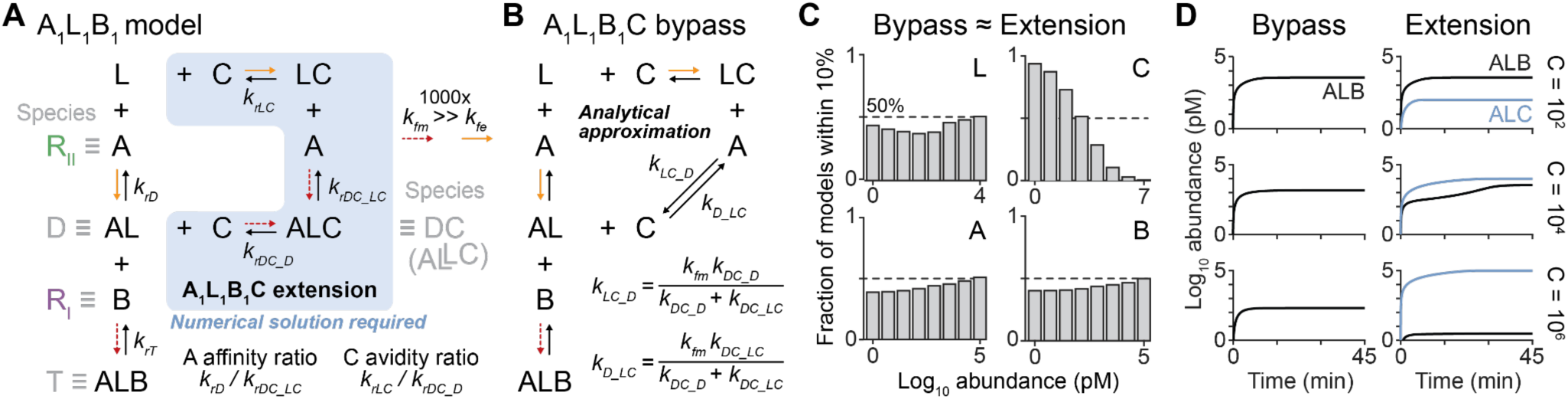
Models of canonical TGFβ signaling must be extended numerically when including coreceptor. (A) Model of a prototypical TGFβ signaling axis containing one ligand (L), one type II receptor (R_II_ ≡ A), and one type I receptor (R_I_ ≡ B) expanded to include one coreceptor (≡ C). Heterodimer (≡ D) and heterotrimer (≡ T) states are annotated by their constituents. The heterodimer-coreceptor (DC) state implies dimeric L.^1,26^ Forward binding rates are assumed constant and faster for on-membrane associations between A, C, and B (*k_fm_*, red dashed) compared to associations with extracellular L (*k_fe_*, orange). Reverse unbinding rates are variable and thus define the relative affinity of A for L with or without C and the bipartite relative avidity of C for L with or without A. Numerical methods are required to solve the extended A_1_L_1_B_1_C model. (B) A pseudo-steady state approximation bypasses ALC, lumps the affinity and avidity ratios of A and C, and yields a constrained algebraic solution for steady state signaling from ALB heterotrimer (STAR Methods). (C) Bypass mimics the extended A_1_L_1_B_1_C model for less than half the log range of C. The fraction of *N* = 8^8^ ∼ 17 million modeled cell instances in which the bypass model falls within 10% of the full A_1_L_1_B_1_C extension is shown as a function of the abundance of L, C, A, or B. (D) One representative cell instance illustrates how the bypassed ALC intermediate (blue) becomes dominant at steady state relative to ALB (black) with moderate-to-high C (in pM). See also Figure S2.

To this core single ligand–receptor model (A_1_L_1_B_1_), we next appended a second ligand-binding path through a TGFβ coreceptor (“C”). For consistency with experiments, we use the binding scheme of TGFBR3—the topology for ENG is identical but the R_II_–R_I_ binding sequence is reversed.^28^ For TGFBR3, ligand binds coreceptor with the same *k_fe_* as R_II_ but unbinds with a separate dissociation rate constant (*k_rLC_*) reflecting its potential difference in affinity. In the model, the LC complex also enters (at a rate proportional to *k_fm_*) a higher-order complex with one R_I_ to form ALC (“DC”), which is reversible according to *k_rDC_LC_*. Structurally, co-receptor occupies the major ligand-binding site for R_II_,^29,30^ but ALC arises in practice from the dimeric nature of TGFβ ligands (Figure 2A).^1,26^ The abstracted DC complex also has an alternative path to dissociate (at a rate proportional to *k_rDC_D_*) yielding coreceptor and the ligand:R_II_ dimer, which is the presumed route to coreceptor-enhanced heterotrimeric signaling.^31,32^ In this extended ligand–receptor–coreceptor model (A_1_L_1_B_1_C), the shared forward rate constants enable coreceptor efficacy to be succinctly summarized by ratios of off-rates for i) the affinity of A to L and ii) the bivalent avidity of C to R_II_-bound L (Figure 2A).

It is tempting to simplify the extended A_1_L_1_B_1_C model with a pseudo-steady state approximation of ALC that bypasses this species through lumped parameters (Figure 2B). The simplification yields a constrained algebraic solution for the steady state of ALB trimer, which is directly comparable to the earlier analytic solution of Antebi et al.^4^ involving multiple TGFβ ligands and receptors (STAR Methods). We compared solutions of the bypass model for ALB trimer with those of the full A_1_L_1_B_1_C extension by interrogating certain model parameters and sampling others. The seven rate parameters and initial conditions for A and B were each sampled at one of eight logarithmically spaced levels to define a “cell instance”. We then solved each cell instance numerically across a range of C and L initial conditions and compared end-time values of ALB with the bypass approximation. Unfortunately, the bypass model was a suitable approximation only for cell instances that involved the lower range of C (Figure 2C). When C was more abundant, the ALC intermediate became a dominant species that deviated from bypass predictions of ALB (Figure 2D). Results were unchanged when the binding cycle of L, C, and A was modeled as a closed thermodynamic system^33^ (Figures S2A and S2B). We concluded that full numerical solutions were required to assess the true contribution of TGFβ coreceptors.

### Coreceptor inhibits most of the signaling landscape for TGFβ heterotrimers

The A_1_L_1_B_1_C simulations were structured to provide an unbiased assessment of TGFβ signaling as a function of coreceptor expression. For each cell instance, we used steady state ALB heterotrimer abundance to read out signaling. One common alternative is R_II_-bound ligand^29-31^ (Table S1), but this was disfavored for agglomerating ALB with AL and ALC, which do not directly phosphorylate R-SMADs (Figure 2A). We set aside nondeterministic instances in which ALB never exceeded ∼100 copies (“n.d.”) and categorized the remainder as increased (“+”), decreased (“–”), or unchanged (“n.c.”) signaling with coreceptor abundance. More complex biphasic (“+,–” or “–,+”) or triphasic (“–,+,–” or “+,–,+”) patterns were similarly counted when they occurred (STAR Methods). The goal was to identify prevailing trends, if any, for coreceptor within the TGFβ signaling architecture (Figure 2A).

For 60% of all cell instances (and 82% of those deterministic), we found that coreceptor was a monotonic inhibitor of heterotrimeric signaling (Figure 3A). Pure enhancement by coreceptor was very rare (1%); it was almost ninefold more common to observe biphasic +,– dependence, whereby ALB enhancements were eventually overtaken by LC and ALC complexes at the highest C. Such biphasic trends are predicted for low-affinity TGFβ ligands by other models,^27^ and we also saw evidence of it (Figure S3A). However, the results here suggest the phenomenon may be more widespread, even without shedding of the coreceptor ectodomain.^34^ The +,– trend generally required the AL affinity for B to be higher than for C (Figures S3B), which is consistent with measurements of TGFBR2:TGFβ1, TGFBR1, and TGFBR3.^29,35^ Compared to –, the +,– trend occurred at high affinity ratios for A, low avidity ratios for C, and low L (Figures 3B–D), possibly explaining why the strongest evidence for this trend is in vivo.^36^

**Figure 3.**
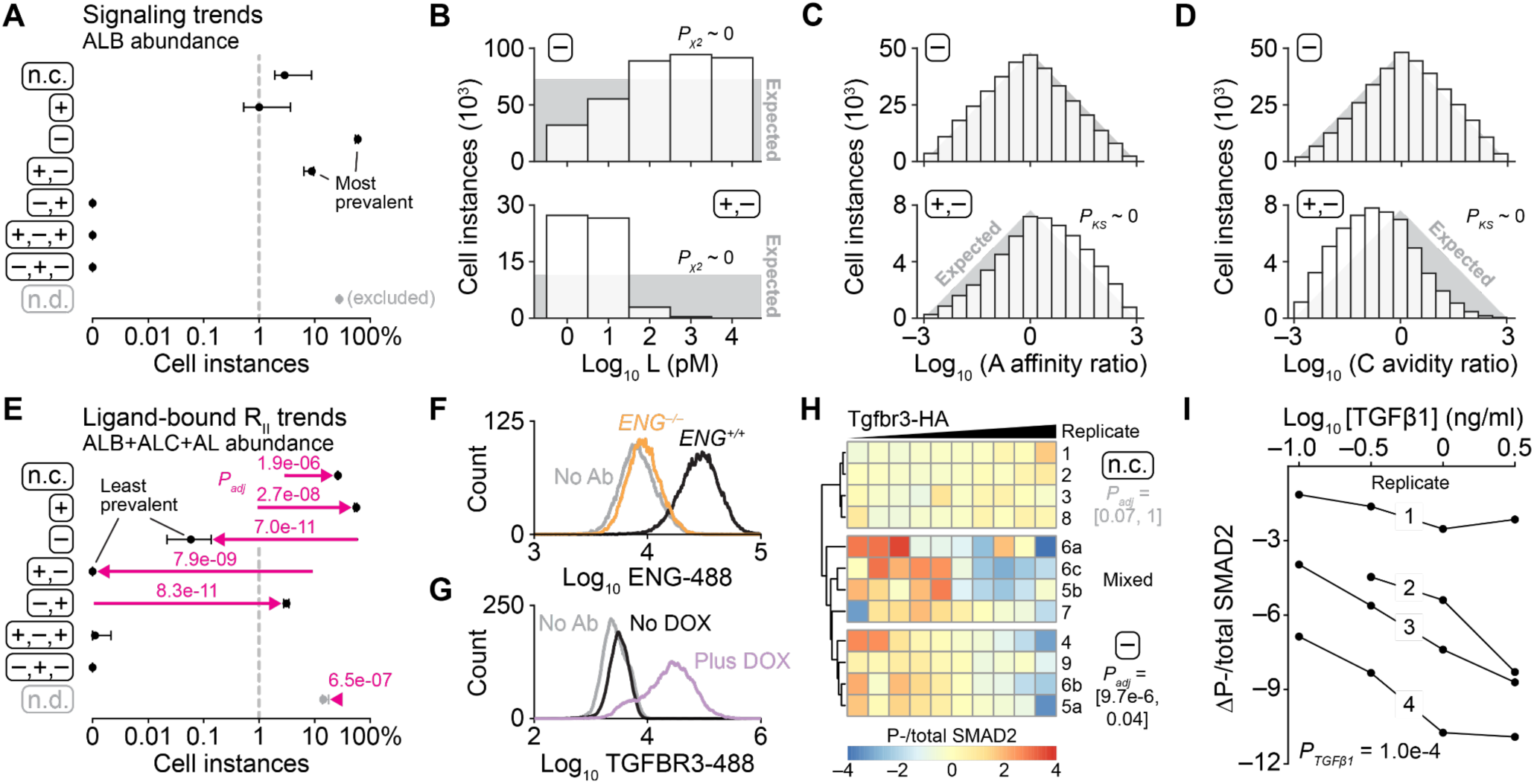
TGFβ signaling architectures are inhibited by coreceptor when ligand or coreceptor abundance is high. (A) Inventory of predicted signaling trends with increasing coreceptor: n.c., less than 10% change between minimum and maximum coreceptor; +, increased signaling; –, decreased signaling; n.d., nondeterministic signaling of fewer than ∼100 copies per cell (excluded). Multiphasic trends of + and – are ordered as a function of increasing coreceptor. Signaling output is proportional to the abundance of ALB (≡ R_II_LR_I_) heterotrimer. (B) Ligand dependence of prevalent coreceptor trends. The – trend is prevalent when L is moderate to high, whereas the +,– trend is most likely when L is low. Differences between the observed (bars) and expected (gray) counts for the discrete L values were assessed by *χ*^2^ test. (C and D) Affinity (C) and avidity (D) dependence of prevalent coreceptor trends. High affinity ratios for ligand by A (≡ R_II_) and low avidity ratios for ligand by C (≡ coreceptor) favor the +,– trend but not the – trend. Differences between the observed (bars) and expected (gray) distribution of affinity or avidity ratios for +,– were assessed by K-S test. See also Figure 2A. (E) Predicted redistribution of coreceptor trends when signaling output is instead taken as proportional to the total abundance of ligand-bound R_II_ (ALB+ALC+AL). Differences between (A) and (E) (magenta) were assessed by paired, arcsine square root transformed *t* test after Šidák correction for multiple-hypothesis testing. (F and G) Flow cytometric confirmation of (F) *ENG* knockout to remove the endogenous coreceptor and (G) doxycycline (DOX)-inducible expression of Tgfbr3-HA in MCF10A cells (STAR Methods). In (G), cells were treated with or without 1 µg/ml DOX for 24 hours and stained with total TGFBR3 antibody to assess the endogenous surface expression without DOX. (H) Variable-yet-recurrent basal signaling trends among cell groups with increasing coreceptor. Engineered *ENG^-/-^* Tgfbr3-HA MCF10A cells were induced with 1 µg/ml DOX for 24 hours, stained for phospho (P-) SMAD2, total SMAD2, Tgfbr3-HA, and DNA, and quantified by flow cytometry. Per-cell P-SMAD2 was normalized to total SMAD2, and cells were stratified within the range of induced Tgfbr3-HA immunoreactivity. Data are shown as the row centered bootstrapped mean ratio of P-SMAD2 and total SMAD2 from *N* = 150–4850 cells per Tgfbr3-HA stratum for 12 independent samples collected and analyzed on nine separate days (numeric prefix). Coreceptor trends were assessed per replicate by ANOVA with Tgfbr3-HA stratum as a continuous factor and Šidák correction for multiple-hypothesis testing. The range of adjusted *P* values (*P_adj_*) is reported in brackets. Replicates in the mixed group co-clustered and had a detectable Tgfbr3-HA effect only when analyzed collectively with replicate as a fixed batch effect (*P_adj_* = 9.0e-03). (I) Ligand stimulation accentuates the – trend of coreceptor. Engineered *ENG^-/-^* Tgfbr3-HA MCF10A cells were induced with 1 µg/ml DOX for 24 hours, stimulated with the indicated concentration TGFβ1 for 30 minutes, stained for P-SMAD2, total SMAD2, Tgfbr3-HA, and DNA, and quantified by flow cytometry as in (H). The – trend magnitude is shown as the difference between the minimum P-/total SMAD2 for a Tgfbr3-HA stratum minus the maximum P-/total SMAD2 for a Tgfbr3-HA stratum (ΔP-total SMAD2) at each TGFβ1 dose for *N* = 4 independent experiments. Trend magnitudes were analyzed by two-way ANOVA with TGFβ1 as a continuous factor and replicate as a fixed batch effect. Significance of TGFβ1 as a continuous factor (*P_TGFβ1_*) is shown. For (A) and (E), data from *N* = 100,000 cell instances are shown as the mean percentage ± range of eight leave-one-out iterations in which one coreceptor abundance was omitted before classification. The dashed line at 1% provides a fixed vertical axis. For (B–D), ∼ 0 is <10^-300^. See also Figure S3.

Other complex patterns were nonexistent for the A_1_L_1_B_1_ case (Figure 3A). By contrast, when the same simulations were re-categorized based on R_II_-bound ligand (ALB + AL + ALC) instead of signaling heterotrimer (ALB), the prevailing trends reversed: – disappeared along with +,– and they were replaced by + and –,+ (Figure 3E). The inconsistent accounting of ALC and fractional observation of +,–trends may explain the preponderance of coreceptor-enhanced readouts in the literature (Table S1).

The A_1_L_1_B_1_C model yielded two actionable predictions: i) coreceptor is almost universally inhibitory when C is highest (combining – and +,– trends of Figure 3A), and ii) inhibitory trends for coreceptor are further enhanced with intermediate-to-high L (Figure 3B). We searched for evidence of these predictions in the MCF10A breast epithelial cell line. MCF10A cells detectably express TGFBR3 and ENG coreceptors,^13,37,38^ and they respond to TGFβ ligands.^13,39-42^ MCF10A cells also express little-to-no β-arrestins,^43^ which mediate endocytosis of ALC complexes.^44^ Starting with a clonal MCF10A derivative,^45^ we isolated TGFBR3 by knocking out ENG with CRISPR-Cas9 and introducing a doxycycline-inducible, HA-tagged murine Tgfbr3 (STAR Methods). After confirming both genetic perturbations by immunoblotting (Figures S3C and S3D) and flow cytometry (Figures 3F and 3G), we sought to measure basal and acute-phase signaling to R-SMADs as a function of induced Tgfbr3-HA. R-SMAD signaling is highly dependent on cell-to-cell protein abundance,^24^ as well as other growth factors added or conditioned into the medium.^46^ We induced Tgfbr3-HA for 20 hours followed by starvation for four hours, which restricted basal R-SMAD signaling to endogenous autocrine ligands (i.e., low-to-moderate L). Then, we concurrently measured phospho-SMAD2, total SMAD2, Tgfbr3-HA, and DNA content (for cell cycle) in single cells by flow cytometry (STAR Methods). The normalized per-cell phospho-/total SMAD2 ratio was binned by decile of HA immunoreactivity to achieve a cell-to-cell variability analysis^47^ of Tgfbr3 function (Figure S3E).

Across unstimulated biological replicates (*N* = 12) we observed insensitivity plus two recurrent Tgfbr3 dose-dependent regimes: – and a mixed group with variable patterns of +,– (Figure 3H). These were the three most-prevalent trends predicted by the A_1_L_1_B_1_C model (Figure 3A). Day-to-day differences may relate to the uniformity of plating^48^ or starvation-induced effects on endogenous TGFBR3 or TGFβ ligands^46^ that are impossible to control fully.^24^ Nevertheless, when we stimulated cells on the same day with increasing TGFβ1 for 30 minutes before processing, we repeatedly observed an increasing magnitude of – dose dependence with co-measured Tgfbr3 (Figure 3I). These experimental results built confidence in the general model architecture and motivated computational expansion to ALB signaling configurations that were more complex.

### R_II_ signaling multiplicity and asymmetry enhance noncanonical dose dependencies for coreceptor Independently of R_I_

In the TGFβ superfamily, multiplicity of receptors enhances ligand discrimination when different receptors bind and signal differently.^4^ We thus considered expansions of the A_1_L_1_B_1_C model that included two R_I_s (A_1_L_1_B_2_C) or two R_II_s (A_2_L_1_B_1_C). A_1_L_1_B_2_C added two additional parameters (*k_rT2_* and the fraction of B allocated to B_2_; Figure 4A), whereas A_2_L_1_B_1_C added four additional parameters (*k_rD2_*, *k_rDC2_D2_*, *k_rDC2_LC_*, and the fraction of A allocated to A_2_; Figure 4B). We accommodated the extra parameters by downsampling the original A_1_L_1_B_1_C space tenfold, preserving the underlying trend distribution (Figure 3A) and then upsampling each new parameter at three or nine levels combinatorially (STAR Methods). Coreceptor trends for the resulting numerical simulations were then summarized under one of two scenarios: i) equal signaling from the two downstream heterotrimers, or maximally unequal signaling with one heterotrimer behaving as a null decoy (Figures 4A and 4B). Scenario (i) measured the impact of the parallel rate processes added to the base A_1_L_1_B_1_C model (Figures 2A, 4A, and 4B). Scenario (ii) was the limiting case for maximal asymmetry between competing heterotrimers; all intermediate cases of signaling should fall in between Scenarios (i) and (ii).

**Figure 4.**
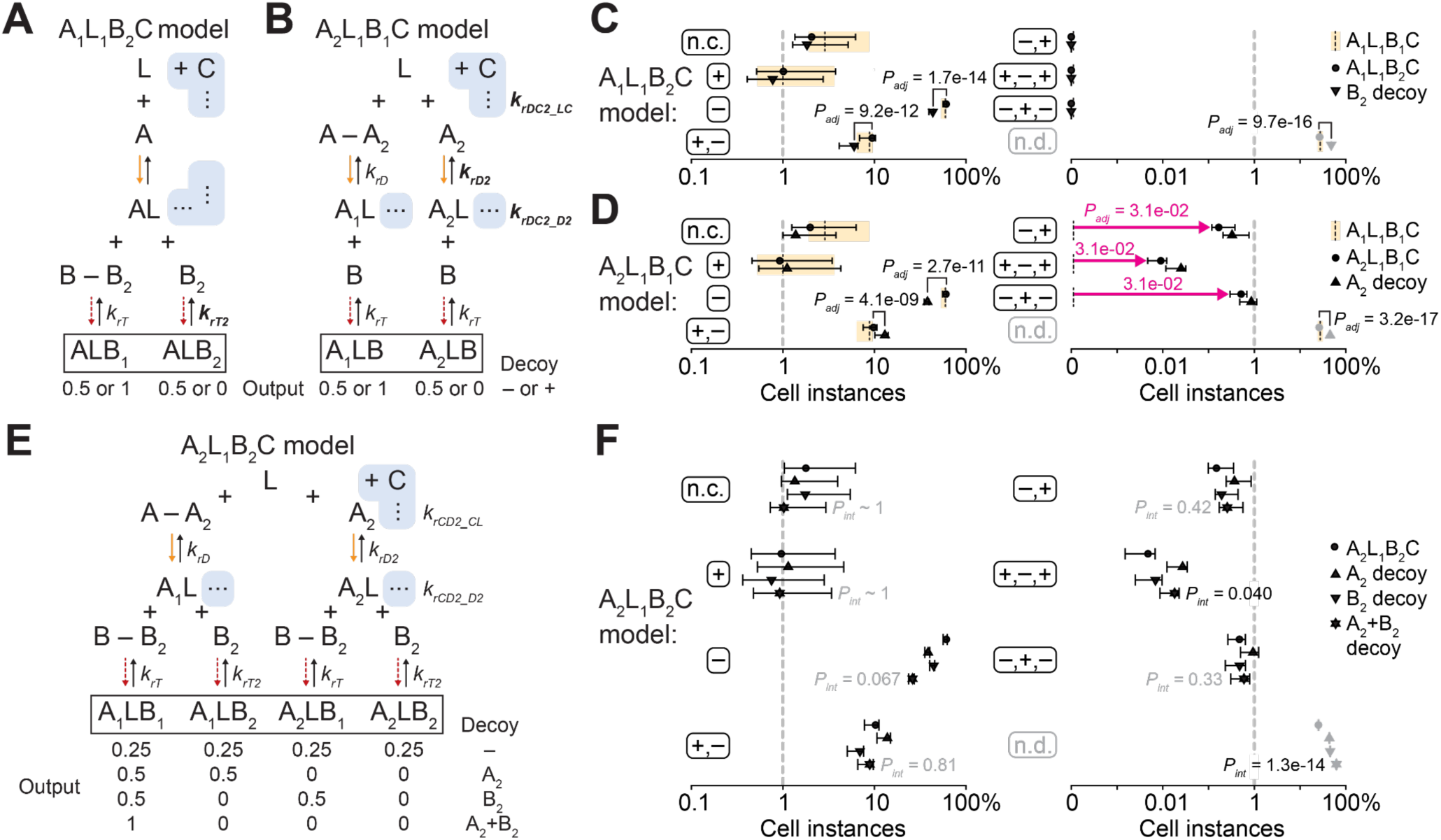
Coreceptor-dependent TGFβ signaling is diversified by competitive multiplicity and asymmetry of R_II_ but not R_I_. (A and B) Expansion of the prototypical TGFβ model from Figure 2A to include (A) two R_I_s, yielding the A_1_L_1_B_2_C model, or (B) two R_II_s, yielding the A_2_L_1_B_1_C model. For both models, ALB heterotrimers contribute to the signaling output equally (0.5 and 0.5) or asymmetrically (1 and 0). The A_1_L_1_B*_2_*C model in (A) adds one sampled parameter (*k_rT2_*), and the A_2_L_1_B_1_C model in (B) adds three (*k_rDC2_LC_*, *k_rD2_*, and *k_rDC2_D2_*) (bolded). The first receptor is mass-balanced to the second [B – B_2_ in (A) and A – A_2_ in (B)] for direct comparisons with the A_1_L_1_B_1_C model. (C and D) Inventory of predicted signaling trends with increasing coreceptor for (C) the A_1_L_1_B*_2_*C model and (D) the A_2_L_1_B_1_C model. Trends are abbreviated as in the A_1_L_1_B_1_C model of Figure 3A, and the predictions of Figure 3A are replotted for reference (dashed bands). The 0.5-and-0.5 output case (circles) is shown alongside the decoy 1-and-0 output case [downward (C) or upward (D) triangles] for comparison. Differences were assessed by one-sided sign-rank test (magenta) or paired, arcsine square root transformed *t* test after Šidák correction for multiple-hypothesis testing. (E) Combination of the A_2_L_1_B_1_C and A_1_L_1_B*_2_*C models to yield A_2_L_1_B*_2_*C. Signaling outputs vary depending on the decoy state of A_2_ and B_2_. (F) The coreceptor-dependent A_2_L_1_B*_2_*C signaling landscape with A_2_+B_2_ decoys (stars) is mostly captured by the superposition of A_2_ decoy (upward triangles) and B_2_ decoy (downward triangles). Differences within trend groups were assessed by multiway ANOVA after arcsine square root transformation. A_2_ decoy and B_2_ decoy were fixed effects, and significance of the interaction term (*P_int_*) was evaluated after Šidák correction for multiple-hypothesis testing For (C), (D), and (F), data from *N* = 100,000 cell instances are shown as the mean percentage ± range of eight leave-one-out iterations in which one coreceptor abundance was omitted before classification. The dashed line at 1% provides a fixed vertical axis.

For R_I_, the impact of multiplicity was intuitive (Figure 4C). Coreceptor trends under Scenario (i) were superimposable with those of A_1_L_1_B_1_C, as expected, and Scenario (ii) increased the frequency of signaling outputs that were nondeterministic and thus excludible. The increase came largely from cell instances that were originally – or +,– for coreceptor (*P* ∼ 0 by hypergeometric test). Therefore, R_I_ multiplicity alone cannot qualitatively alter coreceptor signaling trends aside from silencing them entirely.

By contrast, when R_II_ multiplicity was increased and similarly upsampled, we observed instances of switching between trend classes. The –,+; +,–,+; and –,+,– trends were never detected in A_1_L_1_B_1_C, but together they captured a few percent of cell instances in A_2_L_1_B_1_C (magenta, Figure 4D). 48–54% of these instances were drawn from – trends with a single R_II_, and they all involved low levels of ligand stimulation (Figures S4A and S4B), suggesting physiological relevance. When one R_II_ was considered a decoy, we observed an increase in nondeterministic outputs and a decrease in – trends for coreceptor that were very similar to decoy R_I_; however, for decoy R_II_, +,– trends increased rather than decreased (Figures 4C and 4D). Such shifts occur in cell instances where decoy R_II_ is originally favored—coreceptor then rebalances ligand binding to the signaling R_II_ (+ trend), until it ultimately sequesters ligand from binding R_I_ (+,– trend) (Figure S4C). The results indicate that noncanonical functions for coreceptor like TGFBR3 (i.e., other than –; Figure 3A) are enhanced with multiple R_II_s and signaling asymmetry. The same should occur for coreceptors like ENG when there are multiple R_I_s.

Next, we asked whether multiplicity and signaling asymmetry of R_I_–R_II_ might together have a nonadditive influence on coreceptor function by assembling an A_2_L_1_B_2_C model (Figure 4E). It was not realistic to upsample all six new parameters of A_2_L_1_B_2_C at three levels, because 3^6^ = 729 simulations per condition would require ∼1000-fold downsampling of the base model, likely destabilizing it as a reference. Consequently, we pivoted to a resampling approach, simulating 10^7^ cell instances of the A_2_L_1_B_2_C model and summarizing coreceptor trends when R_I_, R_II_, or both receptors were considered under Scenario (i) or (ii). The resampling approach was first validated by similarly revisiting the A_1_L_1_B_2_C and A_2_L_1_B_1_C models. We found that the salient Scenario (ii) patterns for R_I_ under A_1_L_1_B_2_C and R_II_ under A_2_L_1_B_1_C were also retained for individual decoys under A_2_L_1_B_2_C (Figures 4C, 4D, and 4F). Importantly, nearly all dual-decoy scenarios under A_2_L_1_B_2_C were predictable from the individual additive effects of R_I_ and R_II_ under Scenario (ii) (corrected *P_int_* > 0.05 by arcsine square root-transformed multiway ANOVA; Figure 4F). We conclude that any signaling asymmetries from multiple R_I_s or R_II_s affect overall coreceptor function independently.

### TGFβ ligand decoys are most effective at enhancing + and +,– dose-dependent trends

A natural extension of this study was to consider microenvironments of two ligands with different affinity alongside one or more receptor pairs (A_1_L_2_B_1_C, A_2_L_2_B_1_C, A_1_L_2_B_2_C, and A_2_L_2_B_2_C). However, since each original cell instance included a ligand range, adding a second ligand changed the cell-instance definition such that it was no longer directly comparable to A_1_L_1_B_1_C as before (Figure S5). We circumvented this challenge by resampling as with A_2_L_1_B_2_C but extending to two ligands. We encoded A_1_L_2_B_1_C, A_2_L_2_B_1_C, A_1_L_2_B_2_C, and A_2_L_2_B_2_C separately, resampled 10^5^ cell instances per model, and then compared the relative magnitude of change in coreceptor trends between Scenarios (i) and (ii). Without ligand decoys, there was a considerable decrease in simulations yielding nondeterministic outputs, consistent with the overall increase of ligands in the L_2_ systems (Figures 5A and S5). We noted a corresponding increase in – trends for coreceptor among two-ligand models (Figure 5A), illustrating its default mode of action as an inhibitor of heterodimeric signaling. The A_1_L_2_B_1_C model also gave rise to –,+; +,–,+; and –,+,– trends at prevalences very similar to A_2_L_1_B_1_C (magenta, Figures 4D and 5A); thus, multiplicity upstream (L_2_) or downstream (A_2_) of coreceptor enhances its breadth of dose dependence. With one ligand decoy, there were large relative increases in nondeterministic but also +,– trends (5.1-fold and 2.0-fold; Fig. 5A), proportionally larger than those obtainable with a decoy R_II_ (1.7-fold and 1.3-fold; Fig. 4D). In L_2_ systems, coreceptor may skew both LC and ALC complexes toward productive ALB heterotrimers; in A_2_ systems, coreceptor may only do so through ALC.

**Figure 5.**
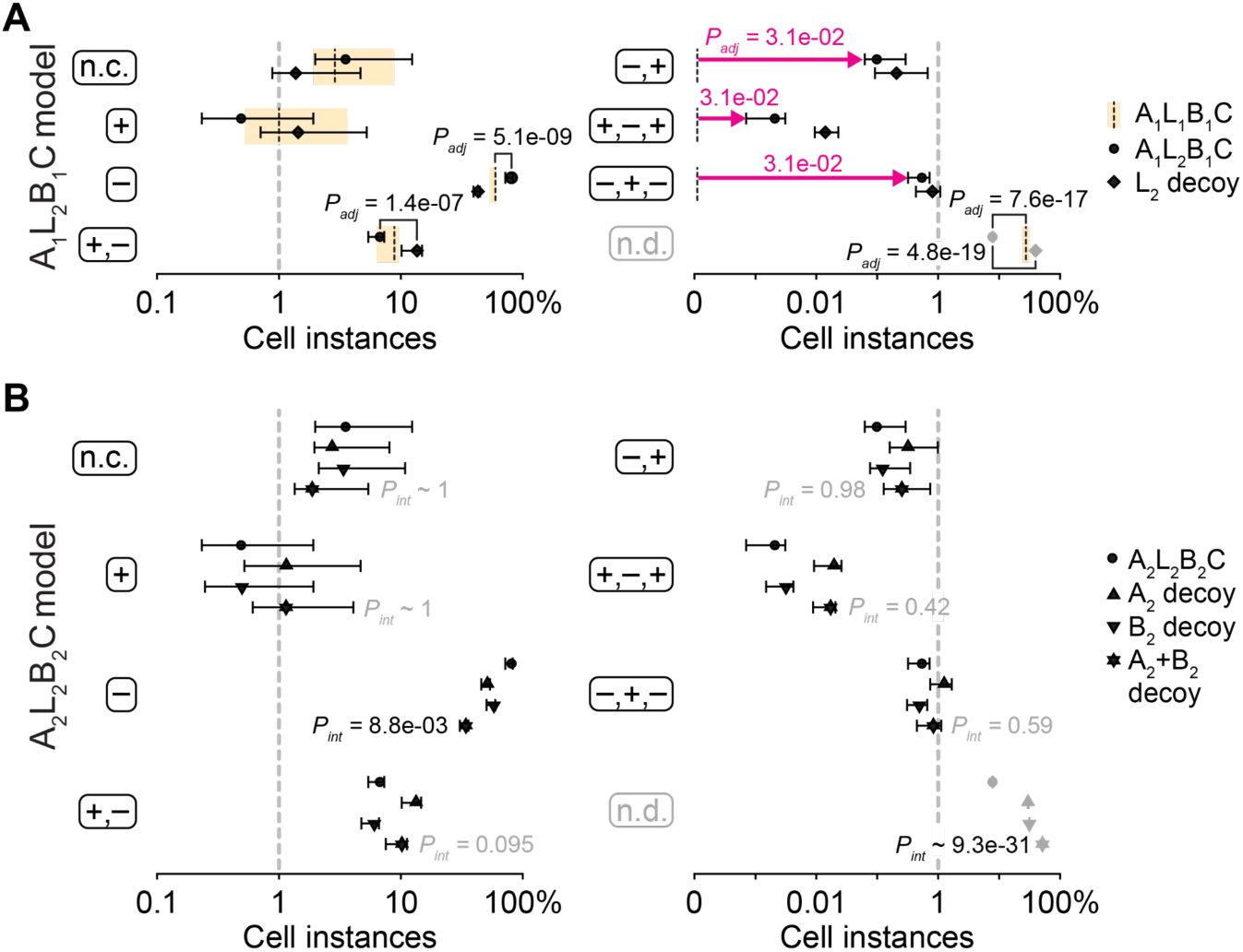
Competitive two-ligand systems promote canonical and non-canonical coreceptor trends independently of the multiplicity or asymmetry of R_I_ or R_II_. (A) Inventory of predicted signaling trends with increasing coreceptor for the A_1_L_2_B_1_C model. Trends are abbreviated as in the A_1_L_1_B_1_C model of Figure 3A, and the predictions of Figure 3A are replotted for reference (dashed band). The 0.5-and-0.5 output case (circles) is shown alongside the decoy 1-and-0 output case (diamonds) for comparison. Differences were assessed by one-sided sign-rank test (magenta) or paired, arcsine square root transformed *t* test after Šidák correction for multiple-hypothesis testing. (B) Two ligands do not appreciably alter the signaling landscape or the independent contributions of A_2_ and B_2_ decoys. Differences within trend groups were assessed by two-way ANOVA after arcsine square root transformation. A_2_ decoy and B_2_ decoy were fixed effects, and significance of the interaction term (*P_int_*) was evaluated after Šidák correction for multiple-hypothesis testing. Data from *N* = 100,000 cell instances are shown as the mean percentage ± range of eight leave-one-out iterations in which one coreceptor abundance was omitted before classification. The dashed line at 1% provides a fixed vertical axis. See also Figure S5.

For the fully combined A_2_L_2_B_2_C model, we revisited the R_I_–R_II_ decoy analysis to see if conclusions changed with two ligands. Models with two receptor decoys yielded somewhat smaller-than-expected decreases in – trends and increases in nondeterministic outputs (Figure 5B). The additional ligand doubles the opportunity to yield productive A_1_LB_1_ signaling heterotrimers and retain the prevailing – trend for coreceptor. We conclude that additional ligands do not substantively impact the R_I_–R_II_ influence on coreceptor function.

### TGFβ coreceptors reconfigure multi-ligand perception in multi-receptor signaling systems

A_2_L_2_B_2_C merited additional analysis, because it was directly compatible with the types of multi-ligand perception described for the A_2_L_2_B_2_ models of Antebi et al.^4^ That work categorized different perception types based on two metrics: a relative ligand strength (RLS) and a ligand interference coefficient (LIC). RLS captures the ratio of maximum signaling from the weaker ligand to that of the stronger ligand (range: 0 to 1). LIC quantifies signaling synergy or antagonism by positively weighing combinatorial signaling above the stronger ligand and negatively weighing combinatorial signaling below the weaker ligand (range: –1 to 1). Together, these metrics capture four A_2_L_2_B_2_ “signaling archetypes”:

i. additive (RLS ∼ 1, LIC ∼ 0): two ligands of comparable signaling strength contribute independently;
ii. ratiometric (RLS ∼ 0, LIC ∼ 0): two ligands of very different signaling strength compete independently for receptors;
iii. imbalance (RLS ∼ 1, LIC < 0): two ligands that each competitively bind to form a weaker ALB signaling heterotrimer; and
iv. balance (RLS ∼ 1, LIC > 0: two ligands that each competitively bind to form a stronger ALB signaling heterotrimer.

Our goal was to examine how signaling archetypes changed when coreceptors were present.

To maintain consistency with Antebi et al.,^4^ we extended the definition of a cell instance by including different heterotrimer signaling efficiencies between 0 (decoy) and 1 (fully active). Upon resampling 10^5^ cell instances, the zero-coreceptor A_2_L_2_B_2_C model generated a bivariate distribution of RLS–LIC values that was qualitatively similar to Antebi et al.^4^ (Figure 6A). The additive archetype was the most prevalent (21% of all instances), followed by ratiometric (4%) and then lower proportions of balance and imbalance (2% each). For the same cell instances, we noted a threshold effect with logarithmically increasing coreceptor, whereby abundances greater than 10^4^ pM caused pronounced changes in both RLS and LIC (Figure 6B). Coreceptor induced different RLS–LIC trajectories depending on the cell instance, but consistently the largest displacements occurred among models with moderate-to-high coreceptor (Figures 6C and 6D, orange). The computational prediction was that multi-ligand perception would be most fragile when coreceptor was very abundant.

**Figure 6.**
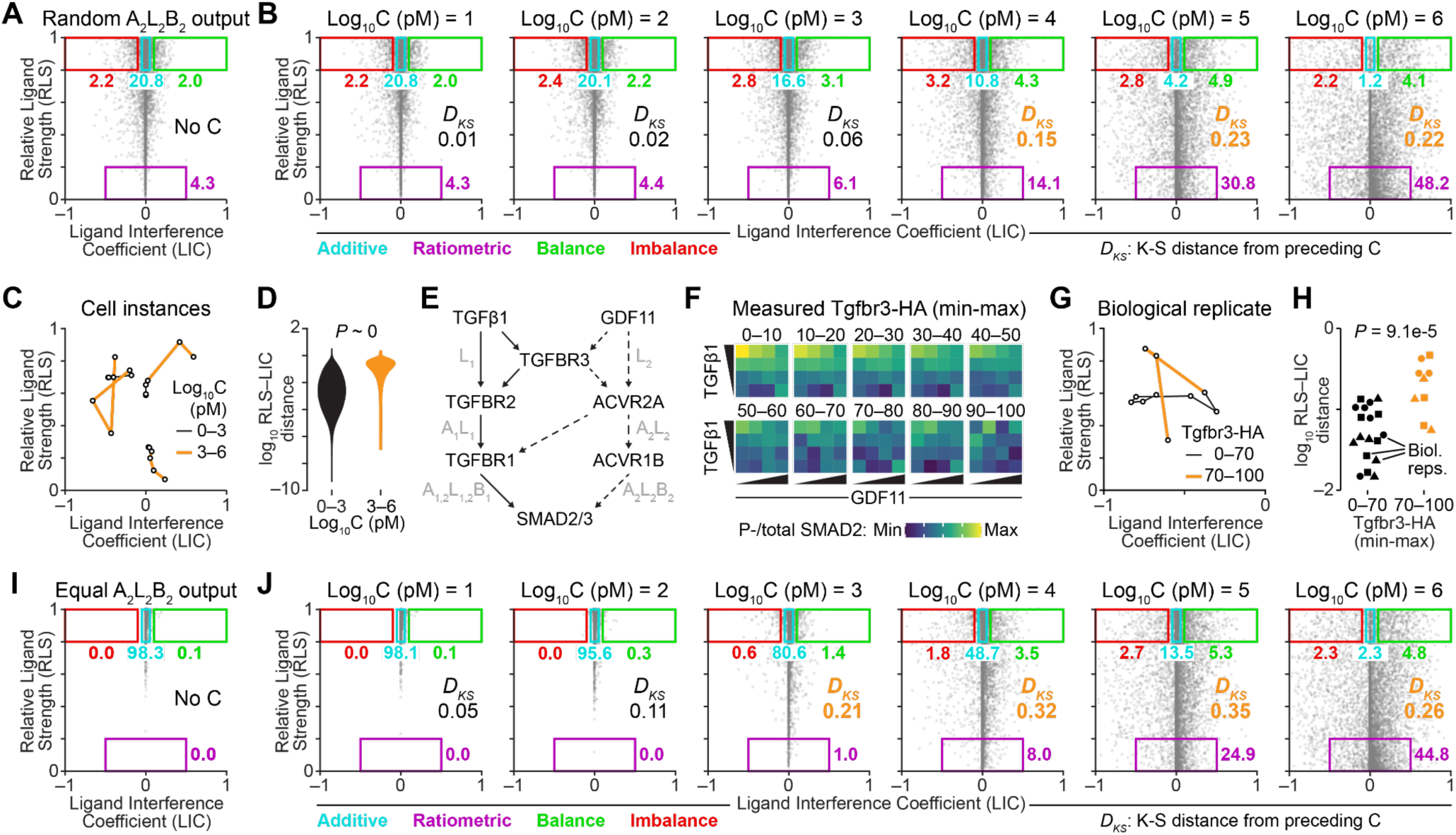
Coreceptor triggers competitive multi-ligand perception by overlapping TGFβ receptors. (A and B) Relative ligand strength (RLS) and ligand interference coefficient (LIC)-based classification of additive (cyan), ratiometric (purple), balance (green), and imbalance (red) signaling archetypes^4^ for two-ligand, two-receptor signaling systems modeled with random signaling efficiencies as output and (A) no coreceptor or (B) increasing coreceptor. (C and D) RLS–LIC deviations are predicted to be most pronounced at moderate-to-high coreceptor. In (C), three illustrative cell instances are shown with edges indicated between cases of low coreceptor [black, log_10_ C (pM) ≤ 3] and moderate-to-high coreceptor [orange, log_10_ C (pM) ≥ 3]. In (D), RLS–LIC Euclidean distances for each category from *N* = 100,000 modeled cell instances are compared by K-S test. (E) TGFβ1 and GDF11 share TGFβ receptors and coreceptor. (F) Engineered *ENG^-/-^* Tgfbr3-HA MCF10A cells were induced with 1 µg/ml DOX for 24 hours, stimulated with TGFβ1 (0, 0.3, 1, 3 ng/ml) and GDF11 (0, 3, 8, 25 ng/ml) for 30 minutes, stained for phospho (P-) SMAD2, total SMAD2, Tgfbr3-HA, and DNA, and quantified by flow cytometry. Per-cell P-SMAD2 was normalized to total SMAD2, and cells were stratified within the range of induced Tgfbr3-HA immunoreactivity. Data are shown as the ratio of bootstrapped means for P-SMAD2 and total SMAD2 from *N* = 150–4850 cells per Tgfbr3-HA stratum. (G) RLS–LIC trajectory for the data in (F) with edges indicated between cases of low coreceptor (black, Tgfbr3-HA = 0–70 percent of the measured range) and moderate-to-high coreceptor (orange, Tgfbr3-HA = 70–100 percent of the measured range). (H) RLS–LIC Euclidean distances between categories from *N* = 3 biological replicates compared by two-way ANOVA with Tgfbr3 category and replicate as factors. (I and J) RLS and LIC-based classification of additive (cyan), ratiometric (purple), balance (green), and imbalance (red) signaling archetypes for two-ligand, two-receptor signaling systems modeled with equal signaling efficiencies as output and (I) no coreceptor or (J) increasing coreceptor. For (A), (B), (I), and (J), the percentages of each signaling archetype from *N* = 100,000 modeled cell instances are reported using the RLS and LIC thresholds of Antebi et al.^4^ The two-dimensional K-S distance (*D_KS_*) of RLS–LIC distributions is compared between increments of coreceptor. Values of *D_KS_* ≥ 0.15 (*P* ∼ 0) are highlighted (orange).

We asked if this prediction was supported by experiments and returned to the engineered MCF10A line with a second ligand alongside TGFβ1. GDF11 is a circulating ligand that signals through different R_II_s (ACVR2A/B) but the same R_I_ (TGFBR1) and R-SMADs as TGFβ1 (Figure 6E).^1,46,49^ At low doses of GDF11, TGFBR3 enhances R-SMAD signaling in MCF10A cells,^41^ suggesting a noncanonical +,– trend that contrasts the purely inhibitory effect of TGFBR3 on TGFβ1 signaling (Figure 3I). We performed two-ligand experiments with a 4x4 matrix of different TGFβ1–GDF11 doses, measured phospho-/total SMAD2 ratio by flow cytometry, and stratified cells by their level of Tgfbr3-HA induction as before (Figure 6F). Cells with >70% of maximum Tgfbr3-HA induction exhibited a markedly different response matrix, with RLS–LIC metrics that deviated erratically compared to cells with lower inductions of Tgfbr3-HA (Figures 6G and 6H, orange). Model and experiment both indicated that multi-ligand perception is redefined by variations in coreceptor abundance (Figures 6D and 6H), which is notable given the large interquartile range of coreceptor transcripts (Figures 1A and 1B).

Last, we repeated the RLS–LIC analysis under the simplifying assumption of equal signaling efficiency from all ALB heterotrimers. In this scenario and without coreceptor, RLS largely remains greater than 0.5, |LIC| generally does not exceed 0.2, and the archetypes are all additive or nearly additive (Figure 6I). By contrast, as coreceptor increased, the models predicted a thresholded dispersion of RLS–LIC values and a re-establishment of all signaling archetypes (Figure 6J). Thus, complete multi-ligand perception does not require different heterotrimer signaling efficiencies if there is a coreceptor that biases ALC complex formation and the handoff to ALB heterotrimers.

## DISCUSSION

This study provides full numerical solutions of coreceptor effects for TGFβ signaling architectures of 1–2 ligands, R_II_s, and R_I_s of varying signal strength. By simplifying forward rates and dimeric ligands, we sampled the remaining parameters realistically and exhaustively to examine the consequence of coreceptor up- or downregulation. TGFβ modeling efforts elsewhere have pointed to the importance of limiting species for the R-SMAD signal;^50^ when more abundant, coreceptors define which complexes become limiting a given ligand dose. Our models predict that for signal transmission to R-SMADs, the net impact is always inhibitory at the highest coreceptor, when ligand and receptor sequester with coreceptor instead of binding to the remaining heterotrimeric partner. This property holds true for rarer affinity–avidity contexts in which coreceptor promotes signaling heterotrimers at low-to-moderate abundances. Such contexts are more achievable with ligands and coreceptor-binding signaling receptors that signal asymmetrically. We saw evidence of both inhibition and promote-then-inhibit in cells engineered to express TGFBR3.

For the multi-ligand case, coreceptor achieves non-additive ligand perception even when all heterotrimers signal equally. Ratiometric perception arises when a – coreceptor trend for one ligand combines with a +,– (or n.c.) trend for a second ligand. The rarer balance and imbalance archetypes emerge from combinations of single ligands with bi- and multiphasic coreceptor trends that become possible with multiple R_II_s. In practice, these situations are somewhat easier to envision than having ALB heterotrimers with the same R_I_ phosphorylate R-SMADs with different efficiency.

We annotated the models according to the current structural understanding of TGFBR3, but the results are equally valid for ENG, which forms a CLB complex (instead of an ALC complex) before heterotrimer formation.^12,28^ ENG overexpression inhibits TGFβ1 signaling,^51^ including in contexts where TGFBR3 overexpression promotes signaling.^52^ The reciprocal binding schemes of TGFBR3 and ENG may explain why R_I_–R_II_ multiplicity is roughly equal at seven and five, respectively. The models here suggest that R_II_ multiplicity and signaling asymmetry enhance signal modulation by TGFBR3, whereas R_I_ asymmetries enhance ENG. By contrast, ligand asymmetries should enhance both, which may explain why there are over 30 TGFβ ligands in mammals.

The findings of this work extend beyond TGFβ signaling. Some heterotypic receptor systems, such as the ErbB family, lack coreceptors and instead use different ligand affinities, receptor abundances, and signaling efficiencies to diversify responses.^53,54^ However, there are other signaling heterotrimers with known coreceptors—CD300f for IL4RA:IL4:IL2RG^55^ and KL for FGFR:FGF23:FGFR^56,57^—that may reflect specific instances of models sampled here. More generally, computational sampling of signaling architectures within biological constraints is powerful, because it defines the predispositions of a binding-reaction topology and then queries whether cells have evolved toward or away from them.

### Limitations of the study

Models were limited to transmembrane coreceptors and did not consider shed proteoforms, which act unambiguously as soluble decoys that inhibit signaling.^34^ Model parameters were varied independently, but the public datasets suggest that some (co)receptors may covary (Figures S1B and S1C). A more consequential limitation of this study is the difficulty we encountered in experimentally testing predictions from the sampled models. The *ENG^-/-^*derivative of MCF10A with inducible Tgfbr3 was a close approximation of a pure A_2_ (ALK5, ACVR1B)–L_2_ (TGFβ1, GDF11)–B_2_ (TGFBR2, ACVR2B)–C (Tgfbr3) context.^13,41,58^ Yet, we struggled with day-to-day variation in Tgfbr3 dosage trends with or without even one added TGFβ ligand. A likely confounder is endogenous GDF11, which is sporadically matured by a sparingly abundant convertase in MCF10A lines.^41,42^ By comparison, multi-ligand experiments combining TGFβ1 and recombinant GDF11 were generally more reliable. Such results further illustrate that coreceptor trends depend on ligand multiplicity and asymmetry.

## Supporting information

Key Resources Table

## RESOURCE AVAILABILITY

### Lead Contact

Further information and requests for resources and reagents should be directed to and will be fulfilled by the Lead Contact, Kevin A. Janes.

### Materials Availability

This study did not generate new unique reagents.

### Data and Code Availability

Code used to generate the results and figures in this paper is available on GitHub (https://github.com/JanesLab/FaresWA_TGFbetaCoR). Source data will be made available in LabArchives (doi to be finalized after revision).

## ACKNOWLEDGMENTS

We thank David Wotton, Eli Zunder, Jeffrey Saucerman, and Matthew Lazzara for critically reading this manuscript and Cameron Griffiths, Piotr Przanowski, Roza Przanowska, Monserrat Gerardo-Ramírez, and Zuping Wang for their review of the figures. This work was supported by grants from the National Institutes of Health (R01-CA214718, U54-CA274499, R01-AI186222 to K.A.J.) and the UVA Comprehensive Cancer Center (PJ04191 to K.A.J.), as well as fellowship support from the National Institutes of Health (T32-CA009109) and National Science Foundation (1842490 to W.A.F.). Data for this manuscript were generated in the University of Virginia Flow Cytometry Core Facility (RRid:SCR_017829), which is partially supported by P30-CA044579.

## AUTHOR CONTRIBUTIONS

Conceptualization: W.A.F., K.A.J.

Data curation: W.A.F.

Formal analysis: W.A.F.

Funding acquisition: W.A.F., K.A.J.

Investigation: W.A.F., K.A.J.

Methodology: W.A.F., K.A.J.

Project administration: K.A.J.

Resources: K.A.J.

Software:

Supervision: K.A.J.

Validation: W.A.F.

Visualization: W.A.F., K.A.J.

Writing – original draft: W.A.F., K.A.J.

Writing – review & editing: W.A.F., K.A.J.

## STAR METHODS

### EXPERIMENTAL MODEL AND STUDY PARTICIPANT DETAILS

#### Genotype-Tissue Expression Dataset

RNA sequencing data were obtained from the Genotype-Tissue Expression project release V8 by downloading ‘GTEx_Analysis_2017-06-05_v8_RNASeQCv1.1.9_gene_tpm.gct.gz’ from their portal. Human donors were included if there were at least 10 tissue samples profiled. TPM data were transformed as log_2_(TPM+1), and the IQR of log_2_(TPM+1) across tissue samples was calculated.

#### Cancer Cell Line Encyclopedia Dataset

RNA sequencing data were obtained from the Cancer Cell Line Encyclopedia (CCLE) by downloading ‘CCLE_RNAseq_rsem_genes_tpm_20180929.txt.gz’ from the Depmap portal. Cell lines were then grouped by tissue type based on CCLE annotation. Tissues with eight or more cancer cell lines were included in downstream analysis. TPM data were transformed as log_2_(TPM+1), and the IQR of log_2_(TPM+1) across tissue types was calculated.

#### The Cancer Genome Atlas Dataset

RNA sequencing data from The Cancer Genome Atlas was acquired by using the TCGAbiolinks R package to identify paired cases with both “Primary Tumor” and “Solid Tissue Normal” sequencing available. Classified tumor types were included if there were at least 10 paired cases profiled. For each case, the log_2_ fold change was calculated as log_2_[(TPM_tumor_+1)/TPM_normal_+1)], and the median log_2_ fold change across cases was calculated for each tumor type.

#### Cell lines

The MCF10A-5E clone (100% match to RRID:CVCL_0598 by STR profiling) and derivatives were cultured as previously described.^45^

### METHOD DETAILS

#### Generalized ALBC extension

We modeled interactions between one TGFBR3-like coreceptor (C) and *n_L_* (= [1, 2]) TGFβ ligands (L), *n_A_* (= [1, 2]) type II receptors (A), and *n_B_* (= [1, 2]) type I receptors (B) as reversible, first-order reactions:

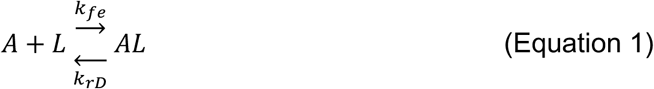

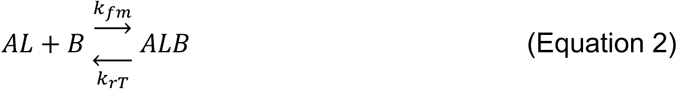

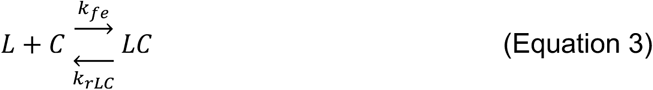

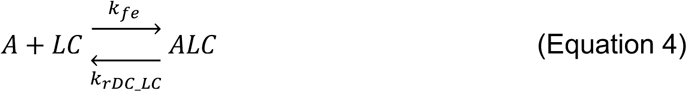

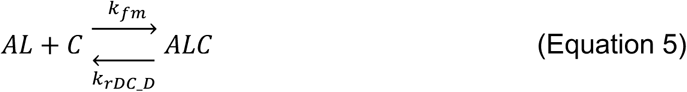

where *k_fe_* is a common association rate for extracellular binding, *k_fm_* is a common association rate for membrane-associated binding, *k_rD_* is the dissociation rate of AL dimer (≡ D), *k_rT_* is the dissociation rate of ALB trimer (≡ T), *k_rLC_* is the dissociation rate of LC complexes, *k_rDC_LC_* is the dissociation rate of ALC complexes (≡ DC) into A and LC, and *k_rDC_D_* is the dissociation rate of ALC complexes (≡ DC) into AL dimer (≡ D) and C.

When expanded to *i* = 1…*n_A_* type II receptors, *j* = 1…*n_L_* ligands, and *k* = 1…*n_B_* type I receptors, Equations 1–5 give rise to the following ordinary differential equations:

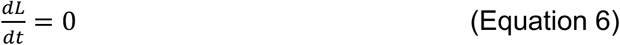

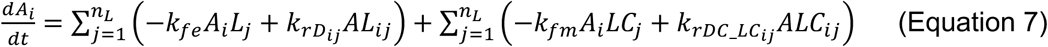

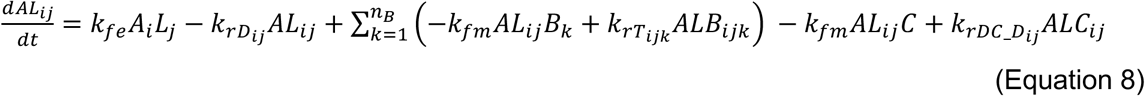

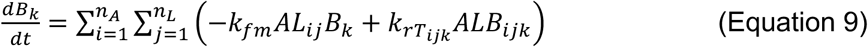

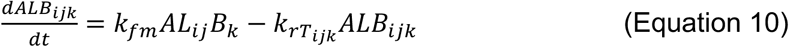

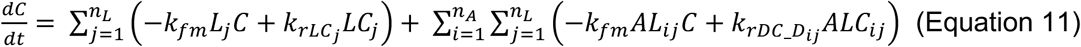

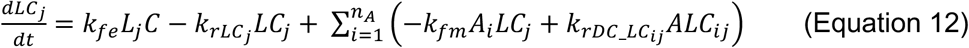

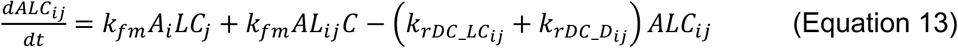

Each ALB heterotrimer was assigned a relative signaling efficiency (*ɛ*, assigned a value of 1 [active] or 0 [decoy]), and total signaling output (*S*) was defined as follows:

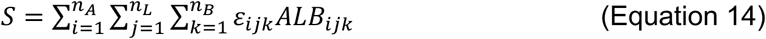

#### ALBC bypass

If ALC is taken as a rare and rapidly equilibrating intermediate, Equations 4 and 5 simplify to:

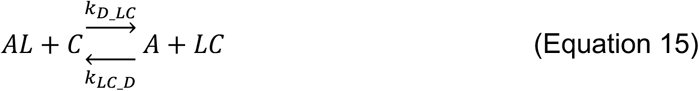

where *k_D_LC_* and *k_LC_D_* are defined by taking a pseudo-steady state approximation of ALC:

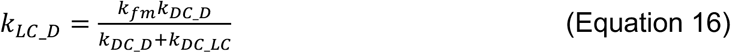

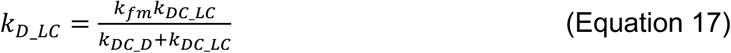

The bypass approximation removes Equation 13 and changes Equations 7–12 as follows:

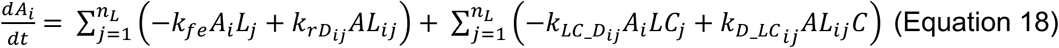

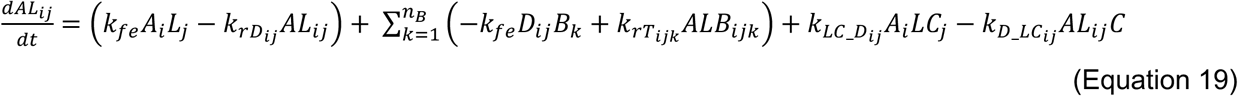

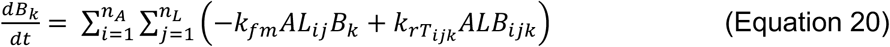

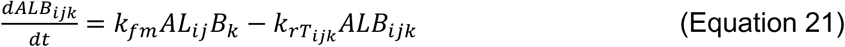

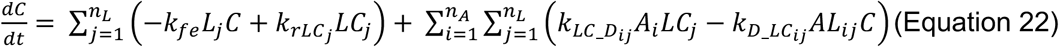

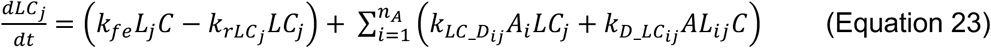

For *n_L_* = 1, *n_A_* = 1, and *n_B_* = 1, if Equations 6 and 17–22 are set equal to zero and solved for steady state solutions by using the following mass balances:

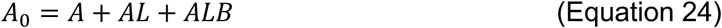

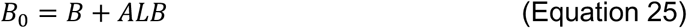

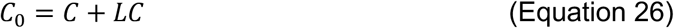

it yields, after rearranging, the following pair of algebraic constraints on steady state ALB heterotrimer (*ALB_SS_*) and LC complexes (LC_SS_):

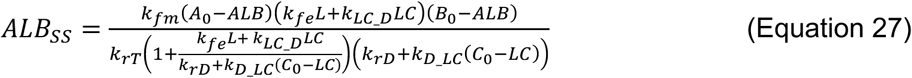

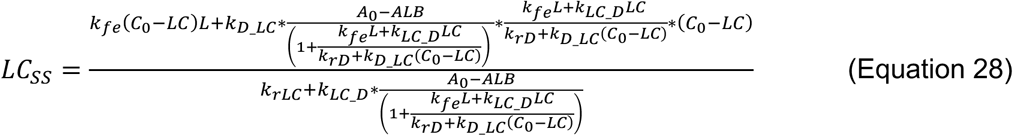

The algebraic constraints generalize for *i* = 1…*n_A_* type II receptors, *j* = 1…*n_L_* ligands, and *k* = 1…*n_B_* type I receptors as follows:

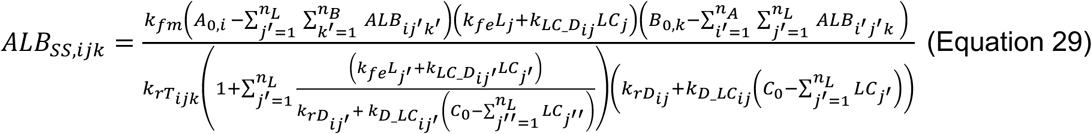

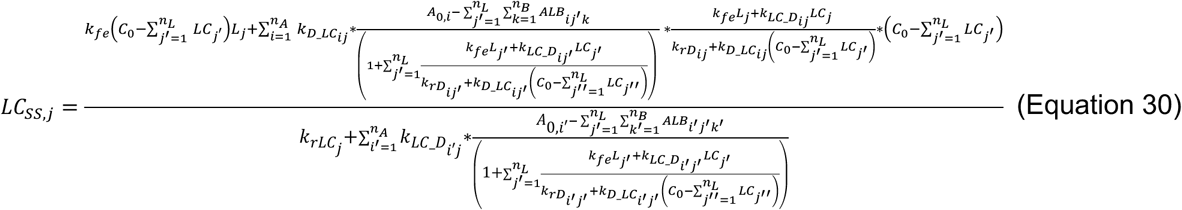

#### Numerical simulations

We used ode15s in MATLAB to solve the systems of differential equations until all time derivatives were less than 1e-2 pM/s.

#### Sampling of Model Parameters

Cell instances were generated by combining fixed ranges for L and C with sampling of other model parameters. When comparing the A_1_L_1_B_1_C extension to the bypass approximation (Fig. 2), we defined eight logarithmically spaced values for L from 1e0–1e4 pM and for C from 1e0–1e7 pM. A and B were discretely sampled at eight logarithmically spaced values from 1e0–1e5 pM. Dissociation rates (*k_rLC_*, *k_rD_*, *k_rDC_LC_*, *k_rDC_D_*, *k_rT_*) were sampled from eight logarithmically spaced values from 1e-5–1e-2 s^-1^. Association rates were kept fixed at *k_fe_* = 10^-7^ pM^-1^s^-1^ and *k_fm_* = 10^-4^ pM^-1^s^-1^. All 8^9^ ∼ 140 million models were numerically solved until steady state and compared pairwise for accuracy of the bypass approximation to within 10% of the full A_1_L_1_B_1_C extension.

For estimating signaling trends with increasing coreceptor (Fig. 3), we used Latin hypercube sampling (LHS) of model parameters other than L and C. We defined six logarithmically spaced values for L from 1e0–1e4 pM and eight values for C: seven logarithmically spaced values from 1e0–1e6 pM plus a 1e-3 pM reference approximating no C. A and B were sampled from 1e2–9e4 pM using a log-uniform distribution. Dissociation rates (*k_rLC_*, *k_rD_*, *k_rDC_LC_*, *k_rDC_D_*, *k_rT_*) were sampled from 1e-5–1e-2 s^-1^ using a log-uniform distribution. Association rates were kept fixed at *k_fe_* = 10^-7^ pM^-1^s^-1^ and *k_fm_* = 10^-4^ pM^-^_1s-1._

For models with type I or II receptor multiplicity, we randomly selected 1% of cell instances from the A_1_L_1_B_1_C model. To model multiple type I receptors, total B was partitioned using nine linearly spaced values from 0.1 to 0.9 into B_1_ and B_2_, and *k_rT2_* was tested at nine logarithmically spaced values from 0.1 to 10 times *k_rT_* for each selected cell instance, yielding 81 perturbations total. To model multiple type II receptors, total A was partitioned 10:90, 50:50, and 90:10 into A_1_ and A_2_, and *k_rD2_*, *k_rT2_*, *k_rDC2_LC_*, and *k_rDC2_D2_* were tested at 0.1, 1, or 10 times the *k_rD_*, *k_rT_*, *k_rDC_LC_*, and *k_rDC_D_* values for each selected cell instance, yielding 243 perturbations total. For models with ligand multiplicity, we examined all 6^2^ = 36 combinations of L_1_ and L_2_ within the prespecified ranges above. Combinations of multiplicity in ligands or type I/II receptors were modeled using 100,000 new cell instances generated by LHS.

#### *ENG* Targeting with CRISPR-Cas9

ENG targeting was performed by Brian Ruis of the UVA Genome Engineering Shared Resource Core. MCF10A-5E cells were electroporated with CleanCap Cas9 mRNA (TriLink, 1 µg/µl stock) and a synthetic sgRNA (100 µM stock) targeting exon 2 of *ENG* (gRNA sequence: ACCACTAGCCAGGTCTCGAA). Roughly one million cells were trypsinized with 0.5% Trypsin (Thermo Fisher Scientific), washed in 1x Dulbecco’s PBS (Thermo Fisher Scientific), and resuspended in 10 µl Neon NxT Genome Editing buffer (Thermo Fisher Scientific) plus 1 µg Cas9 mRNA in 1 µl plus 100 pmol sgRNA, and cells were electroporated using a Neon NxT electroporation system (Thermo Fisher Scientific). Electroporated cells were diluted in 5 ml culture medium, plated immediately in a T25 flask (Corning), and refed after 24 hours. Cells were expanded upon reaching >50% confluence, and editing efficiency was estimated to be 87% by PCR and TIDE analysis.

Residual ENG-positive cells in the edited population were removed by FACS sorting after surface labeling of ENG. Live cells were trypsinized with 0.5% Trypsin (Thermo Fisher) for 5 minutes, gently agitated, and incubated for an additional 3 minutes to ensure complete attachment. Cells were then collected in PBS, spun down at 800 rcf, and washed before blocking in PBS + 1% bovine serum albumin (Fisher BioReagents) (BSA) for 30 minutes at room temperature. After blocking, cells were pelleted and incubated for 30 minutes at room temperature with anti-ENG primary antibody (R&D Systems; 0.25 µg per 10^6^ cells). After staining, cells were incubated for 30 minutes with Alexa Fluor 488-conjugated donkey anti-goat secondary antibody (Thermo Fisher; 1:1000 dilution) before sorting with a Influx cell sorter (BD Biosciences) for the negative population below the first percentile of the *ENG^+/+^*control population. The enriched population was expanded and later confirmed stably *ENG* negative by flow cytometry and immunoblotting.

#### Viral Transduction

Lentivirus was prepared in HEK293T/17 cells (ATCC) by triple transfection of pSLIK-Tgfbr3-HA neo^41^ with psPAX2 + pMD.2G (Addgene) and transduced into *ENG^-/-^*MCF10A-5E cells as described previously.^59^ Transduced cells were selected in MCF10A growth media containing 300 μg/ml G418 until control plates had cleared.

#### Cytokine Stimulation and Cell Isolation

*ENG*^-/-^ Tgfbr3-HA MCF10A-5E cells were seeded at 150,000 cells per well in 12-well plates and cultured for 24 hours. Cells were then treated with 1 μg/ml doxycycline (Sigma-Aldrich) for an additional 24 hours to induce Tgfbr3-HA expression. Following induction, cells were serum-starved for 4 hours. Cells were subsequently treated for 30 minutes with the indicated concentrations of recombinant TGFβ1 (Peprotech) or GDF11 (Peprotech) which were reconstituted according to manufacturer recommendations. After ligand treatment, cells were incubated with 0.5 ml 0.5% trypsin (Thermo Fisher Scientific) for 5 minutes, gently agitated, and incubated for an additional 3 minutes to ensure complete detachment. Plates were transferred to ice, and 1 ml PBS was added to each well. Cell suspensions were collected into pre-chilled 1.6 ml microcentrifuge tubes and centrifuged at 800 rcf for 3 minutes. The supernatant was aspirated, and cell pellets were washed with 1 ml PBS then resuspended in 160 μl TBS and fixed by adding 20 μl room-temperature 37% paraformaldehyde (Thermo Fisher Scientific) (3.7% final concentration in a volume of ∼200 μl) for 15 minutes at room temperature. Fixed samples were centrifuged at 800 rcf and resuspended in 200 μl Tris-buffered saline containing 0.3% Triton X-100 (AquaSolutions) to permeabilize. Samples were stored at 4°C until staining for flow cytometry.

#### Flow Cytometry and Analysis

Fixed samples were pelleted and washed with Tris-buffered saline containing 0.1% Tween-20 (Sigma) (TBS-T). Cells were resuspended in 100 µl 1x Western Block (Roche) diluted in TBS-T, and samples were blocked for 30 minutes on a platform shaker (Jitterbug shaker, speed 6). After blocking, cells were pelleted and incubated for 1 hour at room temperature with primary antibodies in 1x Western Block at the following dilutions: ENG (R&D Systems; 0.25 µg/10^6^ cells), TGFBR3 (R&D System; 0.25 µg per 10^6^ cells), HA (Roche; 0.5 µg/ml), total SMAD2 (Cell Signaling Technology; 1:200 dilution), phospho-SMAD2 (Cell Signaling Technology; 1:400 dilution). After staining, cells were washed three times with 200 μl TBS-T and incubated for 1 hour at room temperature with fluorophore-conjugated secondary antibodies diluted in 1x Western Block at the following dilutions: Alexa Fluor 488-conjugated donkey anti-goat (Thermo Fisher Scientific; 1:1000), Alexa Fluor 488-conjugated goat anti-rat (Thermo Fisher Scientific; 1:1000), Alexa Fluor 647-conjugated goat anti-mouse (Thermo Fisher Scientific; 1:5000), R-Phycoerythrin-conjugated goat anti-rabbit (Jackson ImmunoResearch; 1:400). Cells were washed three times with 200 µl TBS-T, resuspended in 100 µl TBS containing 0.5 μg/ml DAPI (Thermo Fisher Scientific), and analyzed on an Aurora five-laser flow cytometer (Cytek Biosciences). Blank, compensation, and fluorescence-minus-one control samples were included each day.

Scanned samples were gated by forward and side scatter to remove debris and then DAPI fluorescence to isolate nonproliferating cells in G0/G1. Gated cells with HA immunoreactivity below 0.1% or above 99.9% of the observed range were removed as outliers. The remaining cells were partitioned into 10 logarithmically spaced bins and the ratio of phospho-SMAD2 to SMAD2 (after log transformation to account for the detector setup) was calculated for each individual cell.

#### Cell Lysis

For immunoblotting, cells were washed with PBS and lysed in radioimmunoprecipitation buffer plus protease–phosphatase inhibitors: 50 mM Tris-HCl (Sigma) (pH 7.5), 150 mM NaCl (Sigma), 1% (v/v) Triton X-100 (Sigma), 0.5% (w/v) sodium deoxycholate (Sigma), 0.1% (w/v) SDS (Sigma), 5 mM EDTA (Sigma), 10 µg/ml aprotinin (Sigma), 10 µg/ml leupeptin (Sigma), 1 µg/ml pepstatin (Sigma), 1 µg/ml microcystin-LR (Sigma), 200 µM Na_3_VO_4_ (Sigma), and 1 mM PMSF (Sigma). Protein concentrations of clarified extracts were determined with the BCA Protein Assay Kit (Pierce).

#### Immunoblotting

Immunoblotting was performed on 10% polyacrylamide gels with tank transfer to polyvinylidene difluoride membrane and multiplex near-infrared fluorescence detection as described previously.^60^ Primary antibodies were used at the following dilutions: ENG (Thermo Fisher Scientific, 1:1000), TGFBR3 (Cell Signaling Technology, 1:1000), Tubulin (Abcam, 1:20,000), p38 (Santa Cruz Biotechnology, 1:5000), ERK1/2 (Cell Signaling Technology, 1:1000). A dilution of 1:20,000 was used for all secondary antibodies: IRDYE 680LT donkey anti-chicken (LICORbio), IRDYE 800CW goat anti-mouse (LICORbio), IRDYE 800CW goat anti-rabbit (LICORbio), IRDYE 680RD goat anti-rabbit (LICORbio), IRDYE 680LT goat anti-rat (LICORbio).

### QUANTIFICATION AND STATISTICAL ANALYSIS

#### Classification of Coreceptor Trends

We defined signaling output as the steady state concentration of ALB heterotrimers and aggregated multiple heterotrimers as a weighted sum. Heterotrimers were weighed equally unless there was a decoy receptor, in which case all heterotrimers containing the decoy were assigned a weight of zero. The signaling trend with coreceptor for a given cell instance was assigned based on forward differentials of signaling with respect to coreceptor abundance. Differentials were scored if they surpassed a relative change of 10%. Cell instances with no change greater than 10% were considered insensitive to coreceptor and denoted as n.c. Traces that were monotonically increasing or decreasing were denoted + or –, and multiphasic trends were similarly annotated in the order observed with increasing coreceptor. If a cell instance never exceeded 100 pM trimer (∼120 copies for a 2 pL cell volume) for any coreceptor abundance, it was considered nondeterministic and denoted as n.d.

#### Statistics

Methods for hypothesis testing are described in the associated figure caption. For Figure 1, uncertainty in the median values was estimated from 10,000 bootstrap replications and the left-hand percentile of the indicated threshold was used to estimate the *P* value for that threshold. For classifying coreceptor trends (Figures 3–5), bootstrapping of cell instances did not accurately reflect uncertainty in the trend classification. Therefore, we adopted a leave-one-out procedure that omitted one abundance value of C and reclassified to penalize trends that were contingent on only one coreceptor level in a cell instance. The leave-one-out procedure was iterated for all abundance values of C, and conditions were compared recognizing the pairing between leave-one-out subsets. For Figure 6, two-sample, two-dimensional K-S tests^61^ were performed using kstest_2s_2d.m acquired from MATLAB file exchange (Key Resources Table). For the cell-to-cell variability analysis of mean phospho-/total SMAD2 (Figures 3H, 3I, 6F–H, and S3E), 1000 bootstrap replications of the mean were calculated for each HA bin before the indicated ANOVA.

---

**Figure S1.**
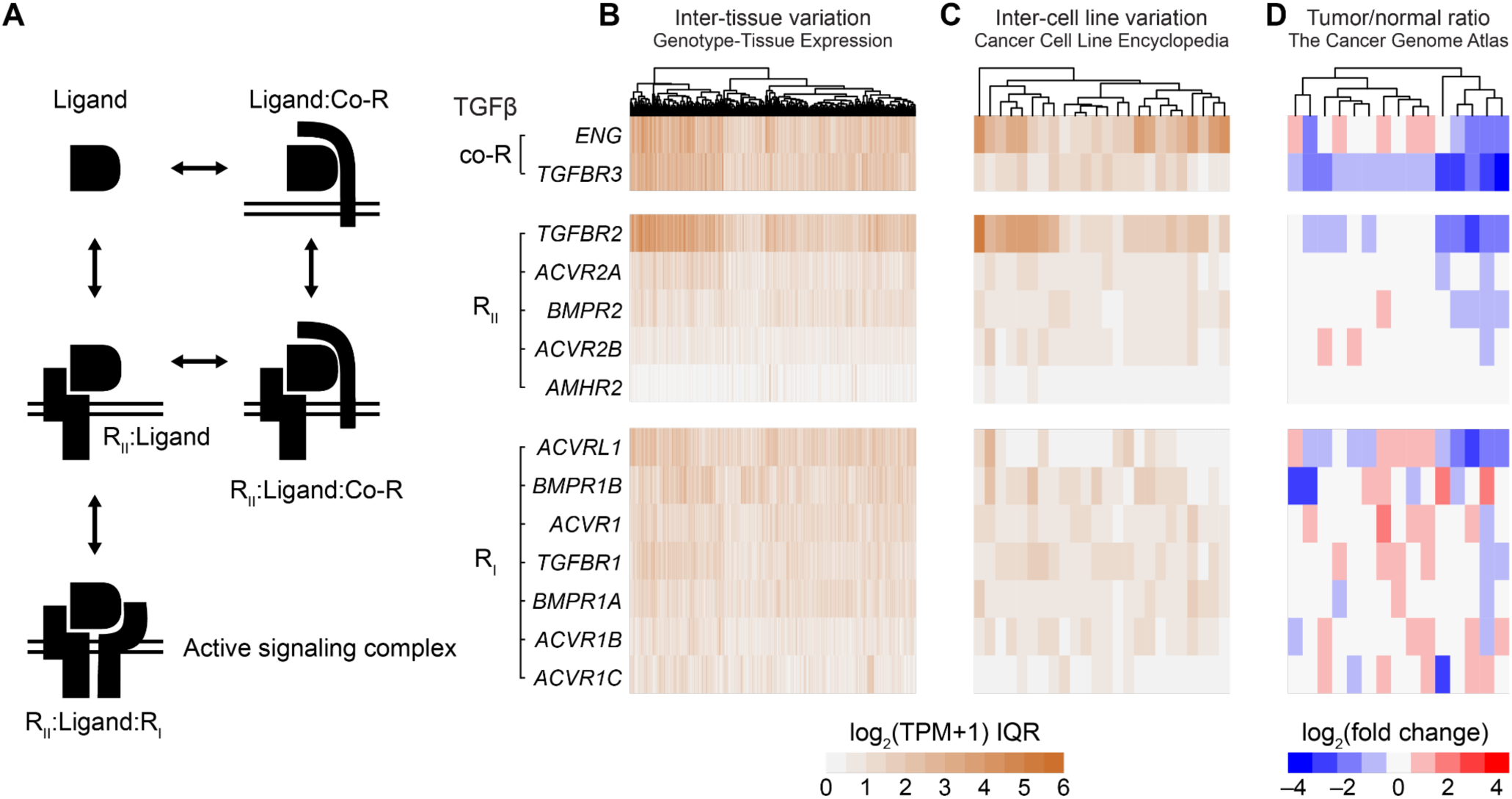
TGFβ (co)receptor variation of individual human donors and cancer types. Related to Figure 1. (A) Schematic relating TGFβ ligand, coreceptor (co-R), type II receptor (R_II_), and type I receptor (R_I_). (B and C) Interquartile range (IQR) of transcript abundance for the indicated TGFβ co-R, R_II_, and R_I_ measured as log_2_ (transcripts per million [TPM] + 1) for (A) *N* = 830 human donors summarized in Figure 1A and (B) the following *N* = 24 cancer types (left to right) summarized in Figure 1B: Soft Tissue, Autonomic Ganglia, Thyroid, Breast, Lung, Prostate, Bone, Stomach, Skin, Biliary Tract, Liver, Esophagus, Upper Aerodigestive Tract, Fibroblast, Large Intestine, Endometrium, Ovary, Central Nervous System, Pancreas, Urinary, Hematopoietic/Lymphoid, Cervix, Kidney, and Pleura. (D) Median log_2_ fold-change estimates of tumor-vs.-normal transcript abundance based on TPM from the following *N* = 15 cancer types (left to right) summarized in Figure 1C: KIRC, KIRP, BLCA, THCA, COAD, PRAD, HNSC, LIHC, ESCA, STAD, BRCA, LUAD, LUSC, KICH, and UCEC.

**Figure S2.**
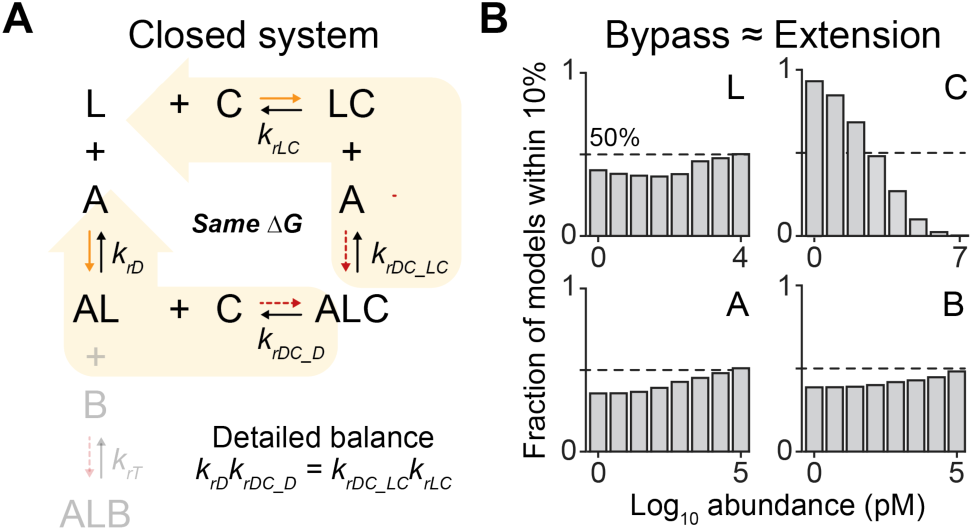
Bypass mimics the extended A_1_L_1_B_1_C model poorly even when the coreceptor loop behaves as a closed system. Related to Figure 2. (A) Considering coreceptor binding and unbinding as a closed system places a thermodynamic constraint relating *k_rD_*, *k_rDC_D_*, *k_rDC_LC_*, and *k_rLC_* by detailed balance.^33^ (B) Same display as in Figure 2C for the 344 subsets of *k_rD_*, *k_rDC_D_*, *k_rDC_LC_*, and *k_rLC_* that conformed to the detailed balance (*N* ∼ 1.4 million modeled cell instances).

**Figure S3.**
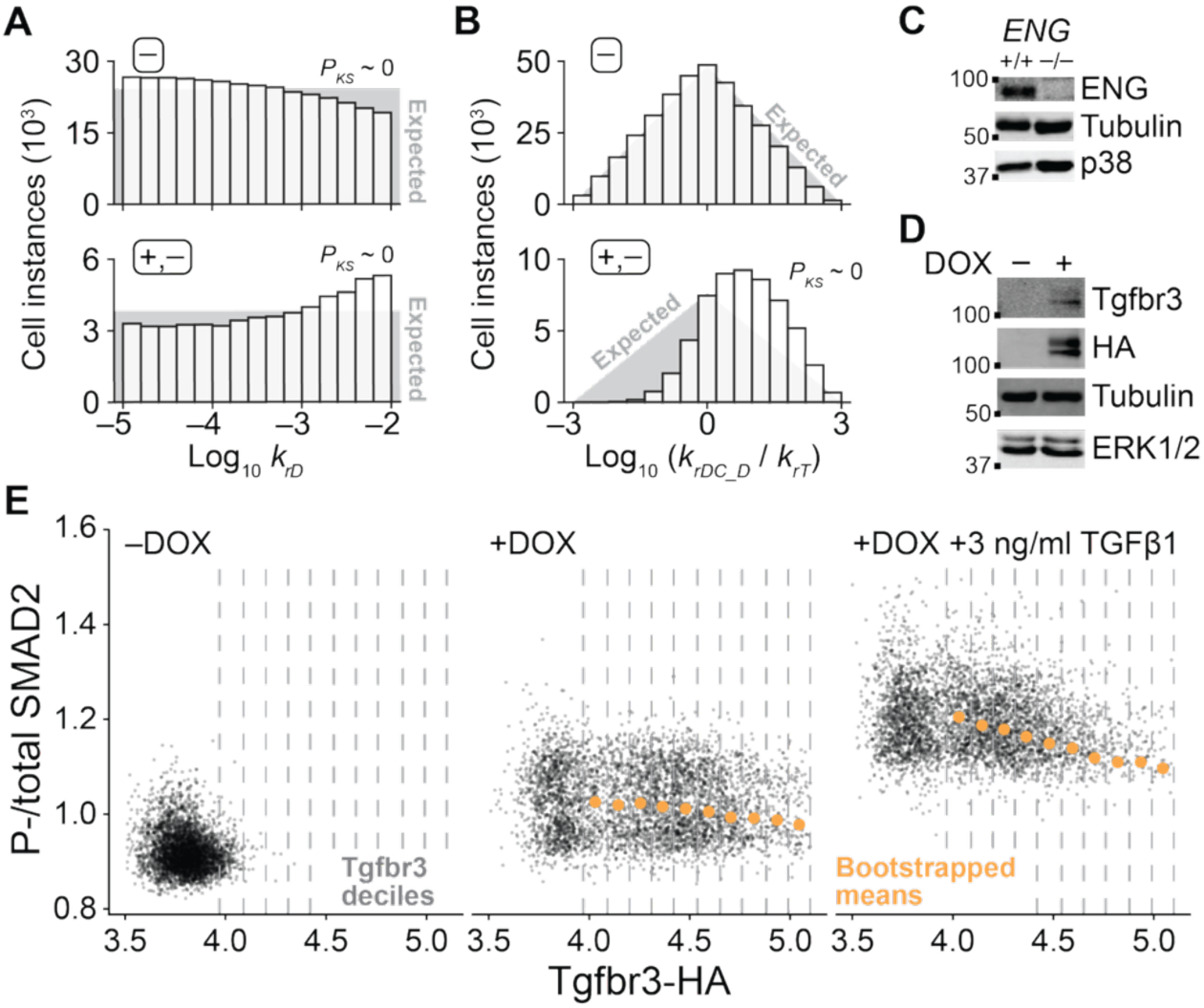
Dependence of prevalent coreceptor trends on additional ligand characteristics and biochemical validation of engineered *ENG^-/-^* Tgfbr3-HA MCF10A cells. Related to Figure 3. (A) Low-affinity ligands with high *k_rD_* for A (≡ R_II_) favor the +,– trend and disfavor the – trend. (B) Ligands with higher affinity for B (≡ R_I_) than for C (≡ coreceptor) favor the +,– trend. (C) Immunoblot of parental MCF10A-5E cells (+/+) and FACS sorted MCF10A-5E *ENG^-/-^* cells (–/–) for ENG with tubulin and p38 used as loading controls. (D) Immunoblot of MCF10A-5E *ENG^-/-^* stably transduced with pSLIK-Tgfbr3-HA, treated with or without 1 µg/ml doxycycline (DOX) for 24 hours, and probed for Tgfbr3 and HA tag with tubulin and ERK1/2 used as loading controls. (E) Representative biplots of Tgfbr3-HA and phospho (P-)/total SMAD2 for *ENG^-/-^* Tgfbr3-HA MCF10A cells treated with or without 1 µg/ml DOX for 24 hours and 3 ng/ml TGFβ1 for 30 minutes. The bootstrapped means for each decile are overlaid. For (A) and (B), differences between the observed (bars) and expected (gray) distributions were assessed by K-S test. ∼ 0 is <10^-300^.

**Figure S4.**
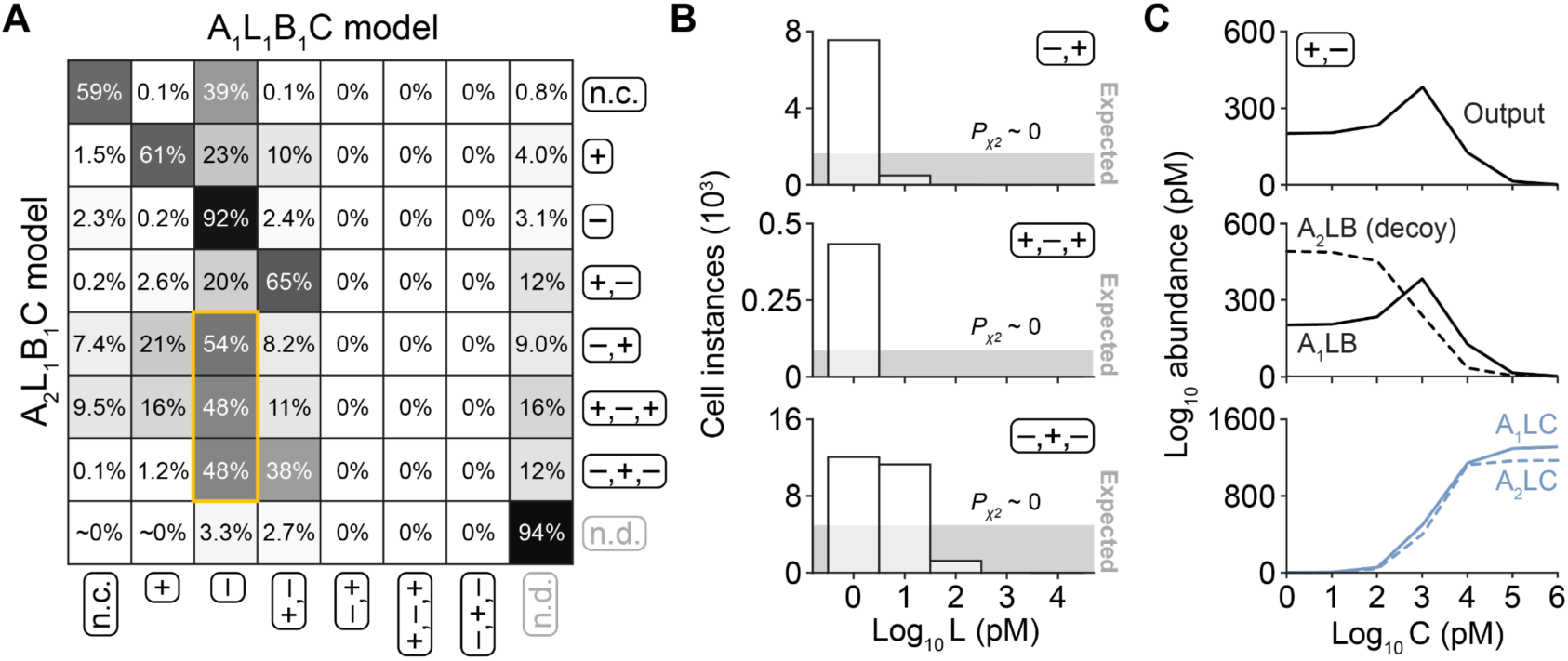
Diverse coreceptor trends with R_II_ multiplicity. Related to Figure 4. (A) Proportion A_2_L_1_B_1_C cell instances in each trend class shared with the A_1_L_1_B_1_C model. The predominant source of the –,+; +,–,+; and –,+,– trends in A_2_L_1_B_1_C are highlighted (yellow box). (B) Ligand dependence of coreceptor trends emerging with R_II_ multiplicity. The –,+ (top); +,–,+ (middle); and –,+,– (bottom) trends are most likely when L is low. Differences between the observed (bars) and expected (gray) counts for the discrete L values were assessed by *χ*^2^ test. (C) One representative cell instance illustrates how the +,– trend emerges with decoy R_II_ as a function of increasing coreceptor. Overall signaling output (top) is shown with the steady state concentration of ALB species (middle) and ALC species (bottom).

**Figure S5.**
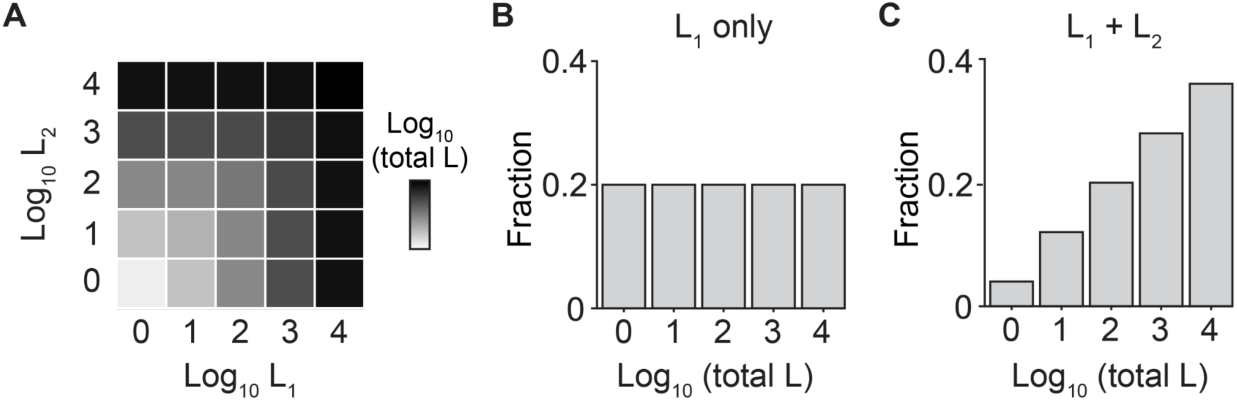
Two-ligand cell instances favor higher total ligand concentrations. Related to Figure 5. (A) Total L displayed as the summation of logarithmically spaced L_1_ and L_2_. (B) Marginal log-uniform density of (A) when considering a single ligand. (C) Overall skewed density of (A) when considering total L.

**Table S1.**
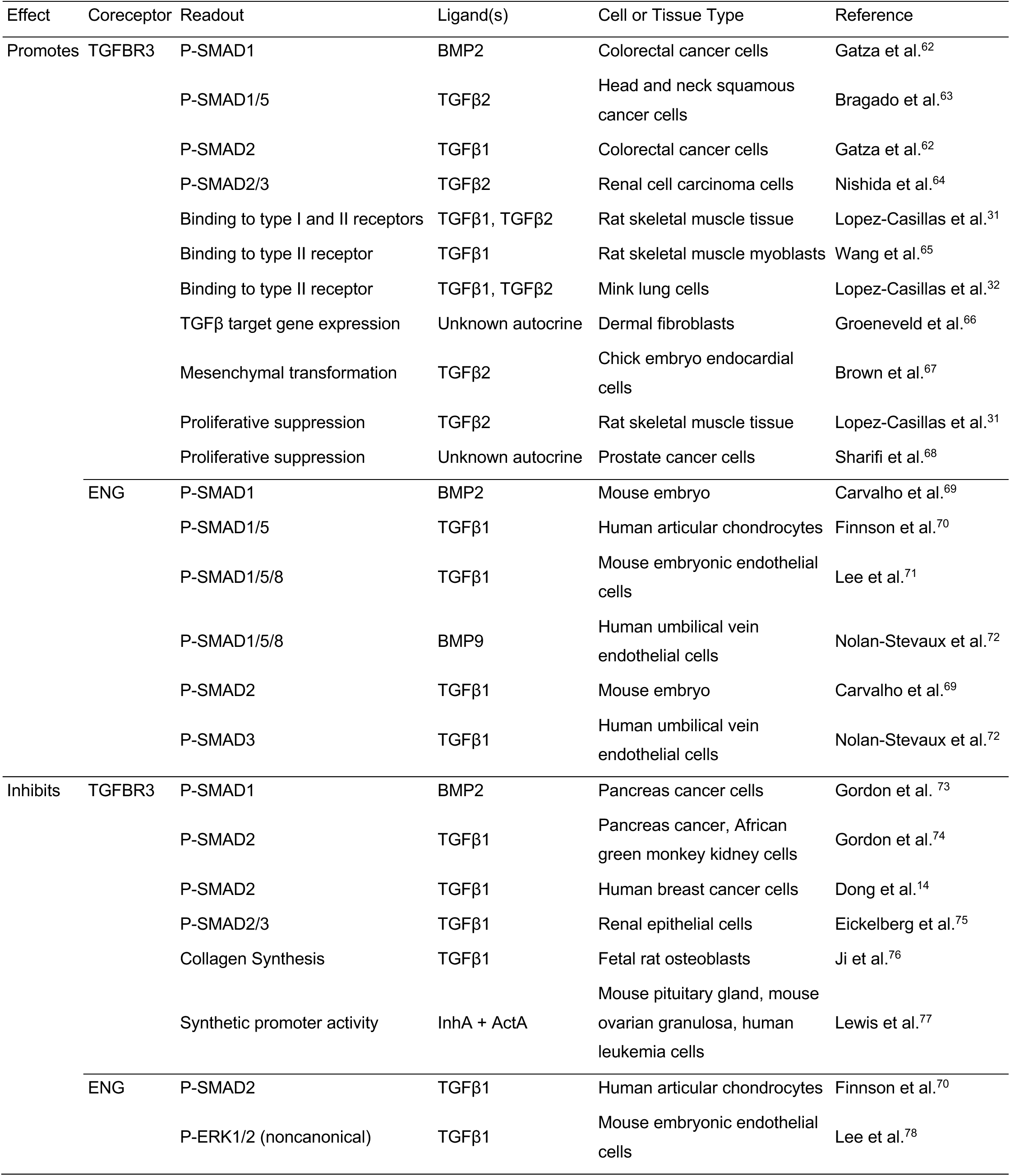
Literature evidence for different coreceptor effects in different cellular contexts. Related to Figure 3.

