## Supplementary material for "TGFβ signaling systems are prone to inhibition and ligand competition by coreceptor": Key Resources Table

| REAGENT or RESOURCE | SOURCE | IDENTIFIER |
| --- | --- | --- |
| Antibodies | | |
| Goat polyclonal anti-ENG | R&D Systems | Cat# AF1097; RRID:AB_354598 |
| Mouse monoclonal anti-ENG | Thermo Fisher | Cat# 14-1057-82; RRID:AB_467417 |
| Rat monoclonal anti-HA | Roche | Cat# 11867423001;  RRID:AB_390918 |
| Rabbit polyclonal anti-p44/42 MAPK (ERK1/2) | Cell Signaling Technology | Cat# 9102; RRID:AB_33074 |
| Rabbit monoclonal anti-phospho-SMAD2 | Cell Signaling Technology | Cat# 18338; RRID:AB_2798798 |
| Rabbit polyclonal anti-p38 | Santa Cruz Biotechnology | Cat# sc-535; RRID:AB_632138 |
| Mouse monoclonal anti-SMAD2 | Cell Signaling Technology | Cat# 3103;  RRID:AB_490816 |
| Goat polyclonal anti-TGFBR3 | R&D Systems | Cat# AF-242-PB;  RRID:AB_354417 |
| Rabbit polyclonal anti-TGFBR3 | Cell Signaling Technology | Cat# 2519; RRID:AB_390707 |
| Chicken polyclonal anti-alpha Tubulin | Abcam | Cat# ab89984; RRID:AB_10672056 |
| IRDYE 680LT donkey anti-chicken | LICORbio | Cat# 926-68028; RRID:AB_10707008 |
| Alexa Fluor 488-conjugated donkey anti-goat | Thermo Fisher | Cat# A11055;  RRID:AB_2534102 |
| Alexa Fluor 647-conjugated goat anti-mouse | Thermo Fisher | Cat# A21236;  RRID:AB_2535805 |
| IRDYE 800CW goat anti-mouse | LICORbio | Cat# 926-32210; RRID:AB_621842 |
| R-Phycoerythrin-conjugated goat anti-rabbit | Jackson ImmunoResearch | Cat# 111-116-144;  RRID:AB_2337985 |
| IRDYE 800CW goat anti-rabbit | LICORbio | Cat# 926-32211; RRID:AB_621843 |
| IRDYE 680RD goat anti-rabbit | LICORbio | Cat# 926-68071; RRID:AB_10956166 |
| Alexa Fluor 488-conjugated goat anti-rat | Thermo Fisher | Cat# A11006; RRID:AB_141373 |
| IRDYE 680LT goat anti-rat | LICORbio | Cat# 926-68029; RRID:AB_10715073 |
| Chemicals, peptides, and recombinant proteins | | |
| Acrylamide ProtoGel (30%) | National Diagnostics | Cat# EC-890 |
| Aprotinin | Sigma | Cat# 10236624001 |
| Bovine serum albumin | Fisher BioReagents | Cat# BP1600 |
| Cholera toxin | Sigma | Cat# C8052 |
| CleanCap Cas9 mRNA | TriLink | Cat# L-7606 |
| DAPI | Thermo Fisher | Cat# 62248 |
| Doxycycline | Sigma | D9891 |
| Dulbecco’s modified Eagle’s medium/F-12 | Gibco | Cat# 11330-032 |
| Dulbecco’s PBS | Thermo Fisher | Cat# 14190094 |
| Eagle's Minimum Essential Medium | ATCC | Cat# 30-2003 |
| Epidermal growth factor | Peprotech | Cat# AF-10015 |
| Ethylenediaminetetraacetic acid (EDTA) | Thermo Fisher | Cat# A10713.0I |
| Fetal bovine serum | HyClone | Cat# SH303396.03 |
| Horse serum | Gibco | Cat# 16050 |
| Hydrocortisone | Sigma | Cat# H0888 |
| Immobilon-FL transfer membrane | Millipore Sigma | Cat# IPFL00010 |
| Insulin | Sigma | Cat# I1882 |
| Leupeptin | Sigma | Cat# 11017101001 |
| Microcystin-LR | Sigma | Cat# 475815 |
| NaCl | Fisher Scientific | Cat# 02-004-047 |
| Paraformaldehyde | Thermo Fisher | Cat# 50980489 |
| Penicillin–streptomycin | Gibco | Cat# 15140 |
| Pepstatin | Sigma | Cat# 10253286001 |
| Phenylmethylsulfonyl fluoride (PMSF) | Sigma | Cat# P7626 |
| Recombinant GDF11 | Peprotech | Cat# 120-11 |
| Recombinant TGFβ1 | Peprotech | Cat# 100-2 |
| Sodium deoxycholate | Sigma | Cat# D6750 |
| Sodium dodecyl sulfate (SDS) | Sigma | Cat# L4509 |
| Sodium orthovanadate (Na3VO4) | Sigma | Cat# S6508 |
| Tris-HCl | Thermo Fisher | Cat# 228030010 |
| Triton X-100 | AquaSolutions | Cat# T9010 |
| Trypsin (0.5%) | Thermo Fisher | Cat# 15400-054 |
| Tween-20 | Sigma | Cat# P9416 |
| Western Blocking Reagent (10x) | Roche | Cat# 11921673001 |
| Critical commercial assays | | |
| BCA Protein Assay Kit | Pierce | Cat# 2322 |
| Neon NxT Electroporation System 10µl Kit | Thermo Fisher | Cat# N1025 |
| Experimental models: Cell lines | | |
| 293T/17 cells | ATCC | Cat# CRL-11268; RRID:CVCL_1926 |
| MCF10A-5E cells | Laboratory of Kevin Janes | Janes et al.^1^ |
| Oligonucleotides | | |
| *ENG* gRNA sequence: ACCACTAGCCAGGTCTCGAA | Synthego | Custom |
| Recombinant DNA | | |
| pMD2.G | Didier Trono Lab | Addgene #12259;  RRID:Addgene_12259 |
| pSLIK Tgfbr3-HA neo | Wang et al.^2^ | Addgene #58508;  RRID:Addgene_58508 |
| psPAX2 | Didier Trono Lab | Addgene: #12260;  RRID:Addgene_12260 |
| Software and algorithms | | |
| MATLAB (version R2025a) | Mathworks | https://www.mathworks.com/downloads/ |
| kstest_2s_2d.m | MATLAB Central File Exchange | https://www.mathworks.com/matlabcentral/fileexchange/38617-kstest_2s_2d-x1-x2-alpha |
| Other | | |
| 10-cm tissue culture plates | Corning | Cat# 430167 |
| 6-well tissue culture plates | Corning | Cat# 3506 |
| Aurora five-laser flow cytometer | Cytek Biosciences | Cat# N7-00003 |
| Influx Cell Sorter | BD Biosciences | Cat# 646500 |
| Jitterbug Microplate Shaker | Boekel Scientific | Cat# 130000 |
| Neon NxT Electroporation System | Thermo Fisher | Cat# NEON1S |
| T25 Flasks | Corning | Cat# 430639 |
